# KANSL1 Deficiency Causes Neuronal Dysfunction by Oxidative Stress-Induced Autophagy

**DOI:** 10.1101/2020.08.07.241257

**Authors:** Katrin Linda, EIly I. Lewerissa, Anouk H. A. Verboven, Michele Gabriele, Monica Frega, Teun M. Klein Gunnewiek, Lynn Devilee, Edda Ulferts, Astrid Oudakker, Chantal Schoenmaker, Hans van Bokhoven, Dirk Schubert, Giuseppe Testa, David A. Koolen, Bert B.A. de Vries, Nael Nadif Kasri

## Abstract

Autophagy is a finely tuned process of programmed degradation and recycling of proteins and cellular components, which is crucial in neuronal function and synaptic integrity. Mounting evidence implicates chromatin remodelling in fine-tuning autophagy pathways. However, this epigenetic regulation is poorly understood in neurons. Here, we investigate the role in autophagy of KANSL1, a member of the nonspecific lethal complex, which acetylates histone H4 on lysine 16 (H4K16ac) to facilitate transcriptional activation. Loss-of-function of KANSL1 is strongly associated with the neurodevelopmental disorder Koolen-de Vries Syndrome (KdVS).

Starting from KANSL1-deficient human induced-pluripotent stem cells, both from KdVS patients and genome-edited lines, we identified superoxide dismutase 1, an antioxidant enzyme, to be significantly decreased, leading to a subsequent increase in oxidative stress and autophagosome accumulation. In KANSL1-deficient neurons, autophagosome accumulation at excitatory synapses resulted in reduced synaptic density, reduced AMPA receptor-mediated transmission and impaired neuronal network activity. Furthermore, we found that increased oxidative stress-mediated autophagosome accumulation leads to increased mTOR activation and decreased lysosome function, further preventing the clearing of autophagosomes. Finally, by pharmacologically reducing oxidative stress, we could rescue the aberrant autophagosome formation as well as synaptic and neuronal network activity in KANSL1-deficient neurons. Our findings thus point towards an important relation between oxidative stress-induced autophagy and synapse function, and demonstrate the importance of H4K16ac-mediated changes in chromatin structure to balance reactive oxygen species- and mTOR-dependent autophagy.

## Introduction

Autophagy is a well-conserved, cellular process controlling the degradation and recycling of proteins, which is essential for cells to maintain protein homeostasis. Three major types of autophagy have been identified: chaperone-mediated autophagy, micro-autophagy, and, most extensively studied, macro-autophagy (hereafter referred to as autophagy). There is emerging evidence for an important physiological role of autophagy in neuronal health and function^1,2^. In neurons, autophagy plays a pivotal role in synaptic plasticity and memory formation^3^. Within the most prominent neurodegenerative disorders, like Alzheimer’s disease (AD), Parkinson’s (PD), and Huntington’s disease, autophagy is protective against neurodegeneration^4,5^. Disrupted autophagy showed to promote neurodegeneration in those disorders^6–8^. During development autophagy is required for synaptic pruning. Impaired autophagy due to over-activation of the mammalian target of rapamycin (mTOR) leads to reduced synaptic pruning and subsequent increased spine density, which has been observed in patients with autism spectrum disorder ^9^. Further, to enable activity-dependent modifications in synaptic strength and efficacy, high protein turnover is required in which autophagy has been proposed to play a role, at the pre- and postsynapse, by controlling the number of synaptic vesicles and ionotropic GABA and glutamate receptors^10–12^.

In recent years, compelling evidence has revealed a crucial role for tight transcriptional regulation of autophagy related (ATG) genes. Several epigenetic mechanisms, including histone modifications, are essential for the transcriptional control of these ATG genes and thus autophagy ^3^. Under nutrient-rich conditions the dimethylation of histone H3 at lysine residue 9 (H3K9me2) represses ATG gene expression in HeLa and primary human T-cells^13^. Similarly, H3K27me3 represses transcription of several negative regulators of mTOR, ensuring high mTOR activity in order to suppress autophagy^14^. Furthermore, ATG gene expression is regulated by the acetylation of histone H4 at lysine residue 16 (H4K16ac), which is mediated by the acetyltransferase hMOF (also KAT8/ MYST1)^15^. While H4K16ac generally ensures active ATG gene expression, autophagy induction is followed by a reduction in H4K16ac and decreased ATG gene expression^15^. This negative feedback-loop prevents prolonged autophagy and subsequent cell death, indicating how essential balanced autophagy is for cell survival, but also function.

The histone acetyltransferase hMOF works within the non-specific lethal (NSL) complex where KANSL1 functions as a scaffold protein^16^. Heterozygous loss of *KANSL1* has been identified as the genetic cause for Koolen-de Vries syndrome (KdVS), formerly known as 17q21.31 micro-deletion syndrome^17–19^. This multisystemic disorder encompasses mild to moderate ID, developmental delay, epilepsy, distinct facial features, congenital malformations, and friendly behavior. In HeLa cells, KANSL1 has been implicated in microtubule nucleation and stabilization during mitosis^20^ and, more recently, also in mitochondrial respiration^21^. In mouse and *Drosophila* genetic manipulations of *KANSL1* have implicated KANSL1/ hMOF signaling in short-term memory formation and social behavior^18,22^. In the KANSL1-deficient mouse model, the behavioral deficits in mice were accompanied with impaired synaptic transmission^22^. Accordingly, changes in gene expression in these mice have been linked to synapse function, development, as well as neurogenesis^22^. However, the molecular and cellular processes through which KANSL1 affects neuronal function, in particular in human neuronal networks, remains unaddressed. The function of KANSL1 as scaffold of the NSL complex and the essential role of H4K16ac in autophagy regulation suggest that deregulated autophagy might underlie the neuronal deficits in KdVS.

Starting from KANSL1-deficient human induced-pluripotent stem cells (iPSCs), both from KdVS patients and genome-edited lines, we identified a previously unrecognized mechanism in which loss of KANSL1 resulted in autophagosome accumulation due to increased oxidative stress. In KdVS patient-derived neurons, a disbalance between mTOR- and oxidative stress-mediated autophagy resulted in reduced synaptic function and network activity, which could be rescued by lowering the amount of reactive oxygen species (ROS). Our findings establish ROS-mediated autophagy as a contributing factor to KdVS presenting a new avenue for autophagy modulation in neurons and possible therapeutic intervention.

## Results

### Heterozygous loss of KANSL1 leads to accumulation of autophagosomes in iPSCs

To examine the role of KANSL1 in autophagy regulation, we generated an isogenic heterozygous KANSL1-deficient iPSC line (CRISPR_1_, Fig. 1A). We used CRISPR/Cas9 genome editing, in a healthy control iPSC line (C_1_, mother from KdVS individual, Supplementary Figure 1B-D), to induce a heterozygous frameshift mutation in exon 2, leading to a premature stop-codon. Additionally, we reprogrammed fibroblasts from a KdVS patient with small deletions in exon 2 of *KANSL1* (KdVS_1_), leading to a frameshift and premature stop-codon. KdVS_1_ was generated from the daughter of C_1_. The shared genetic background of C_1_, CRISPR_1_ and KdVS_1_ allowed us to examine phenotypes caused by heterozygous loss of *KANSL1*. To ensure that our findings are common in KdVS patient-derived cells and not specific to the shared genetic background, we additionally compared lines with an independent genetic background. A second KdVS patient iPSC line (KdVS_2_) was generated from a female patient with a mutation in Exon 2 that is similar to the mutation in KdVS_1_. The third KdVS iPSC line (KdVS_3_) was generated from a patient with a 580-kb heterozygous deletion at chromosome 17q21.31 encompassing five known protein-coding genes (*CRHR1, SPPL2C, MAPT, STH*, and *KANSL1)*. Notably, there is no distinction between KdVS patients with *KANSL1* mutations versus 17q21.31 deletion at the clinical level^19^. Finally, we obtained a gender-matched (female) external control iPSC line (C_2_), which we used for comparison with the KdVS patient lines. All selected clones were positive for pluripotency markers (OCT4, TRA-1-81, and SSEA4) (characterization of iPSCs is illustrated in Supplementary Figure 1A). As expected, Western blot, immunocytochemistry and quantitative real-time PCR (qPCR) analysis showed reduced *KANSL1* expression in all patient lines, as well as the CRISPR-edited line (Figure 1B, Supplementary Figure 1E, 2A). Loss of KANSL1, as scaffold protein of the NSL complex, however, did not lead to a general reduction in H4K16ac (Supplementary Figure 1F), which is in alignment with recent observations of *Nsl1* knockdown in mouse embryonic stem cells^23^.

**Figure 1.**
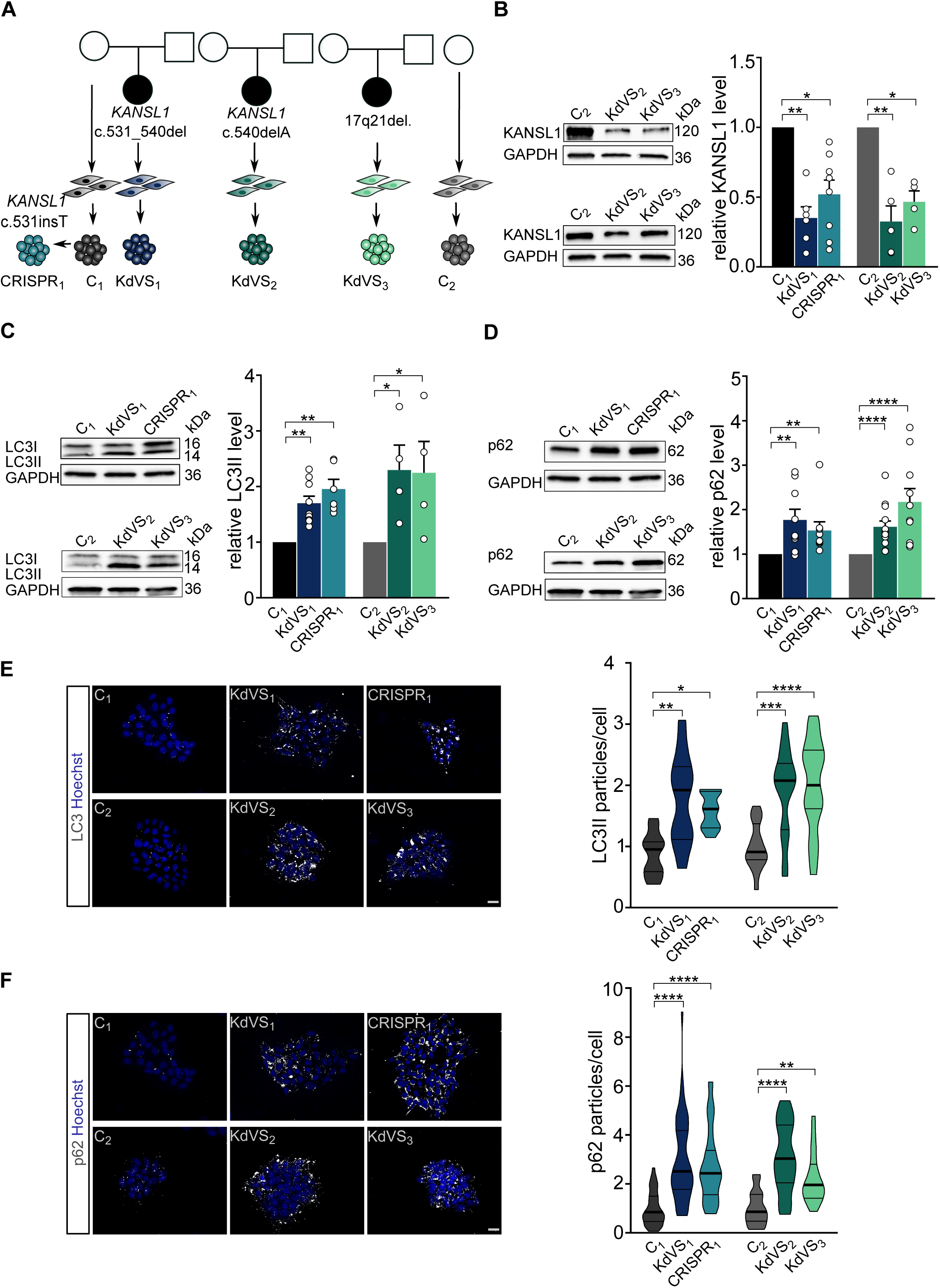
Autophagosome accumulation in KdVS patient derived iPSCs. **A)** Schematic overview of control and KdVS iPSC lines used in this study. **B)** Quantification of KANSL1 protein levels relative to respective control line and example Western blots for all iPSC lines. N = 8 for C_1_; n = 6 for KdVS_1_; n = 8 for CRISPR_1_; n = 5 for C_2_; n = 5 for KdVS_2_; n = 4 for KdVS_3_. **C)** Example Western blots and quantification for LC3II protein levels normalized to control. n = 9 for C_1_; n = 9 for KdVS_1_; n = 6 for CRISPR_1_; n = 4 for C_2_; KdVS_2_; and KdVS_3_. **D)** Example Western blots and quantification for p62 protein levels relative to the respective control line. n = 12 for C_1_; n = 9 for KdVS_1_; n = 9 for CRISPR_1_; n = 13 for C_2_; n = 13 for KdVS_2_; n = 10 for KdVS_3_. **E)** Representative images and particle quantification for iPSC colonies stained for LC3. Scale bar = 20 µm. n = 10 for C_1_; n = 10 for KdVS_1_; n = 11 for CRISPR_1_; n = 16 for C_2_; n = 17 for KdVS_2_; n = 17 for KdVS_3_. Particle quantification was normalized to respective control line. One-way ANOVA and Sidak’s multiple comparison test was used to determine statistically significant differences. **F)** Representative images and particle quantification for iPSC colonies stained for p62. Scale bar = 20 µm. n = 39 for C_1_; n = 45 for KdVS_1_; n = 35 for CRISPR_1_; n = 16 for C_2_; n = 17 for KdVS_2_; n = 19 for KdVS_3_. Data presented in this figure was collected in at least 3 independent experiments. Statistically significant differences were tested through Kruskal-Wallis and Dunn’s multiple comparison test, if not mentioned differently. \**P*□<□0.05, \*\**P*□<□0.01, \*\*\**P*□<□0.005, \*\*\*\**P*□<□0.0001.

To investigate the role of KANSL1 in autophagy we measured autophagosomal marker proteins p62 and LC3I/II in control and KANSL1-deficient iPSCs under basal conditions. Western blot analysis showed increased p62 as well as LC3II levels (Figure 1 C-D) in all KANSL1-deficient iPSCs. We corroborated these results by immunocytochemistry, in which we observed increased numbers of p62 and LC3 particles in all KANSL1-deficient iPSCs, when compared to their respective controls (Figure 1 E-F). LC3II is a well-characterized marker for autophagosomes^24,25^. During acute activation LC3I is lipidated to LC3II and then locates to the double membrane of autophagosomes^26^. Next to elevated LC3II levels, we found increased expression of *WIPI2*, encoding a protein that binds to the early forming autophagosomes, where it is involved in LC3 lipidation^27^ (Supplementary Figure 2B), strongly suggesting increased autophagosome formation in KANSL1-deficient iPSCs. P62 is an “adaptor” protein that interacts with LC3II when bound to targets like protein aggregates or mitochondria, to promote their selective uptake and degradation^28^. Upon fusion with lysosomes, lysosomal enzymes degrade autophagosomal content, including p62, while LC3II at the outer membrane remains stable (reviewed by Glick et al.)^29^. The fact that we observed increased levels of LC3II and p62 under basal conditions, even in the absence of inhibitors of lysosomal proteolysis, indicates that autophagosomes accumulate in KANSL1-deficient iPSCs.

**Figure 2.**
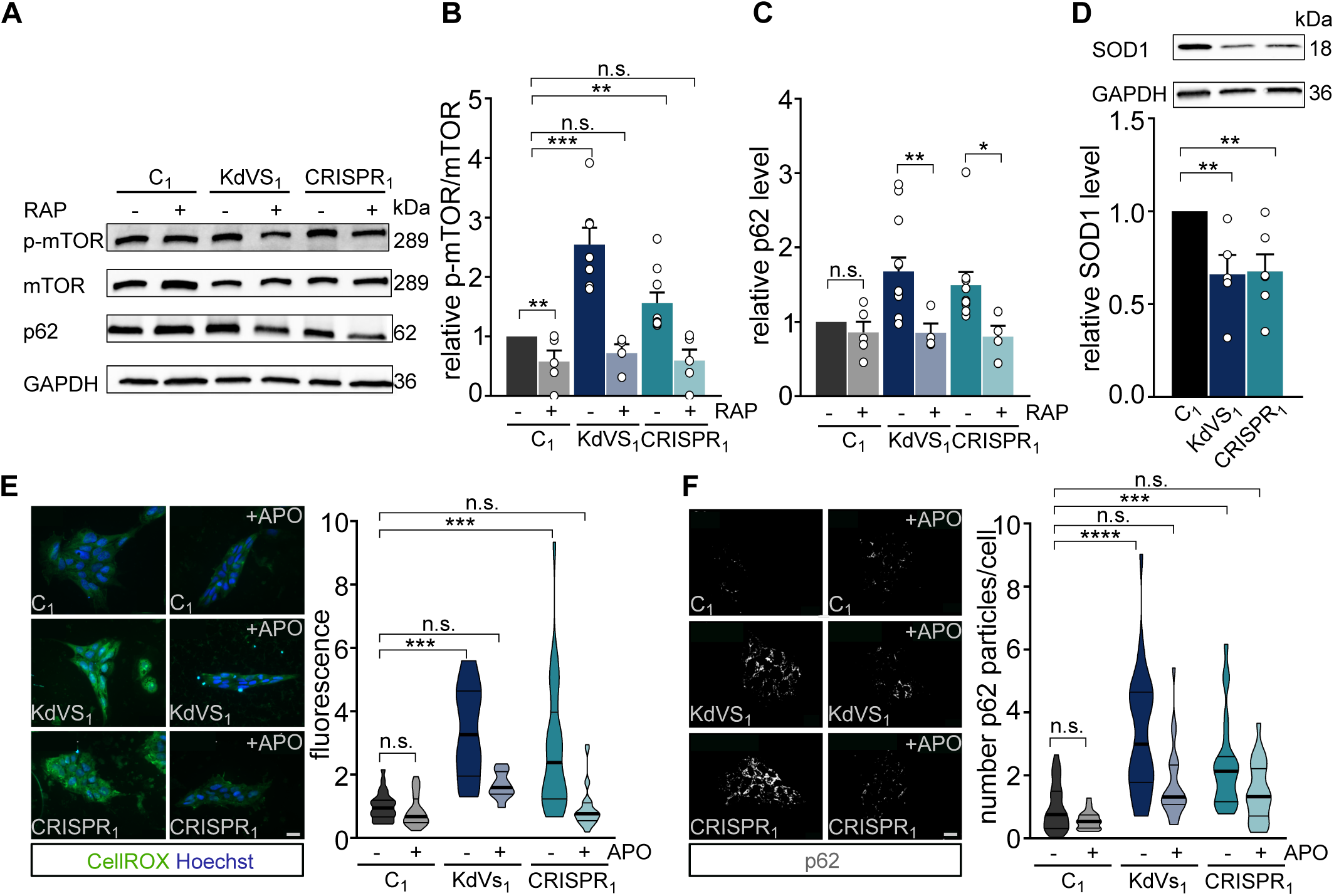
Increased oxidative stress causes autophagosome accumulation. **A)** Representative Western blots for p-mTOR, mTOR and p62 with and without Rapamycin (RAP) treatment. RAP samples were incubated with 200 µM Rapamycin for 10 minutes, followed by 2h without Rapamycin before cell lysis. **B)** Quantification of p-mTOR/ mTOR ratio. n = 9 for C_1_; n = 5 for C_1_ +RAP; n = 7 for KdVS_1_; n = 5 for KdVS_1_ +RAP; n = 9 for CRISPR_1_; n = 5 for CRISPR_1_ +RAP. Protein levels were normalized to untreated control. Treatment efficiency was assessed by means of unpaired t test between untreated and RAP treated samples of C_1_. **C)** Quantification of p62. n = 8 for C_1_; n = 5 for C_1_ +RAP; n = 12 for KdVS_1_; n = 4 for KdVS_1_ +RAP; n = 10 for CRISPR_1_; n = 4 for CRISPR_1_ + RAP. Protein levels were normalized to untreated control. **D)** Representative Western blots for SOD1 and SOD1 protein quantification. Protein levels were normalized to control. n = 6 for C_1_; n = 5 for KdVS_1_; n = 6 for CRISPR_1_. **E)** Representative images of iPSC colonies stained with CellROX and fluorescence quantification. iPSCs were either untreated or treated overnight (o/n) with 100 µM apocynin (APO). n = 26 for C_1_; n = 23 for C_1_+APO; n = 12 for KdVS_1_; n = 14 for KdVS_1_+APO; n = 26 for CRISPR_1_; n = 28 for CRISPR_1_+APO. Fluorescence levels were normalized to untreated control. Scale bar = 20 µm. **F)** Representative images of iPSC colonies stained for p62 and particle analysis. iPSCs were either untreated or treated o/n with 100 µM APO. n = 27 for C_1_; n = 24 for C_1_+APO; n = 33 for KdVS_1_; n = 22 for KdVS_1_+APO; n = 23 for CRISPR_1_; n = 23 for CRISPR_1_+APO. Fluorescence levels were normalized to untreated control. Scale bar = 20 µm. Data presented in this figure was collected in at least 3 independent experiments. Statistically significant differences were determined by means of Kruskal Wallis and Dunn’s multiple comparison test. \**P*□<□0.05, \*\**P*□<□0.01, \*\*\**P*□<□0.005, \*\*\*\**P*□<□0.0001.

### Autophagosome accumulation is caused by increased oxidative stress

The mTOR pathway is a central regulator of autophagy, controlling multiple aspects including initiation, course, and termination of the autophagy process (Supplementary Figure 2D). Upon activation, the mTOR complex 1 (mTORC1) represses autophagy by phosphorylating ULK1 and thereby repressing its’ kinase function within the early steps of autophagosome biogenesis^30–32^. To test whether autophagosome accumulation in KANSL1-deficient iPSCs is caused by reduced activity of mTORC1 we measured p-mTOR, which is the phosphorylated, activated form of mTOR. Surprisingly, we found an increased p-mTOR/ mTOR ratio in KANSL1-deficient iPSCs, indicating increased mTOR activity and thus reduced mTOR mediated autophagy (Figure 2A-B). Increased phosphorylation of ULK1 at serine residue-757^33^ in KANSL1-deficient iPSCs further confirmed a reduction in mTOR mediated autophagy (Supplementary Figure 2C). To test whether the mTOR-associated autophagy pathway is intact in KANSL1-deficient iPSCs we treated the cells with rapamycin, a compound that activates autophagy through mTOR inhibition. Rapamycin reduced p-mTOR levels, as well as p62 protein levels in KANSL1-deficient iPSCs, normalizing the levels between genotypes. This indicates that autolysosome formation and subsequent protein break-down are functional upon rapamycin-induced autophagy (Figure 2A, C). Accumulation of autophagosomes was, therefore, neither mediated by a decrease in mTOR signaling, which even showed to be increased, nor due to impaired lysosomal function in KANSL1-deficient iPSCs.

To identify mTOR-independent molecular mechanisms that can lead to increased autophagosome formation/ accumulation, we considered chromatin-immunoprecipitation sequencing data on differentially activated promoters (H3K4me3) in the KdVS mouse model^22^. Gene ontology (GO) analysis identified changed promoter activation of genes within oxidoreductase and mitochondria pathways^22^, suggesting increased oxidative stress in KdVS mice. We performed qPCRs for a set of H4K16ac regulated ATG genes^15^, that showed differentially activated promoters in the KdVS mouse model. We identified superoxide dismutase 1 (SOD1), an antioxidant enzyme that reduces superoxide, to be about one third lower expressed in KdVS patient derived iPSCs (Supplementary Figure 2A). Western blot analysis further confirmed lowered SOD1 protein levels (Figure 2D). Notably, the expression of the closely related paralog SOD2, which functions in the mitochondria, was unaffected (Supplementary Figure 2A). Reduced expression of antioxidant enzymes, like SOD1, leads to less efficient ROS neutralization and increased oxidative stress. Since increased ROS can activate autophagy in an mTOR-independent manner^34^, we hypothesized that oxidative stress could be causal to the elevated autophagosome formation in KANSL1-deficient cells. To test this hypothesis, we directly measured ROS in control and KANSL1-deficient iPSCs, using a fluorogenic, cell-permeable dye, that exhibits green fluorescence upon oxidation by ROS. We found higher fluorescence levels in KANSL1-deficient cells (Figure 2E), suggesting increased oxidative stress. Because loss-of-function of different NSL complex proteins, including KANSL1, has previously been associated with reduced mitochondrial function in non-neuronal cells^21^, we assessed mitochondrial respiration utilizing Seahorse assay. However, our results indicate that the oxygen consumption rate was not affected in KANSL1-deficient iPSCs (Supplementary Figure 3), suggesting that increased oxidative stress is not occurring due to reduced mitochondrial function.

**Figure 3.**
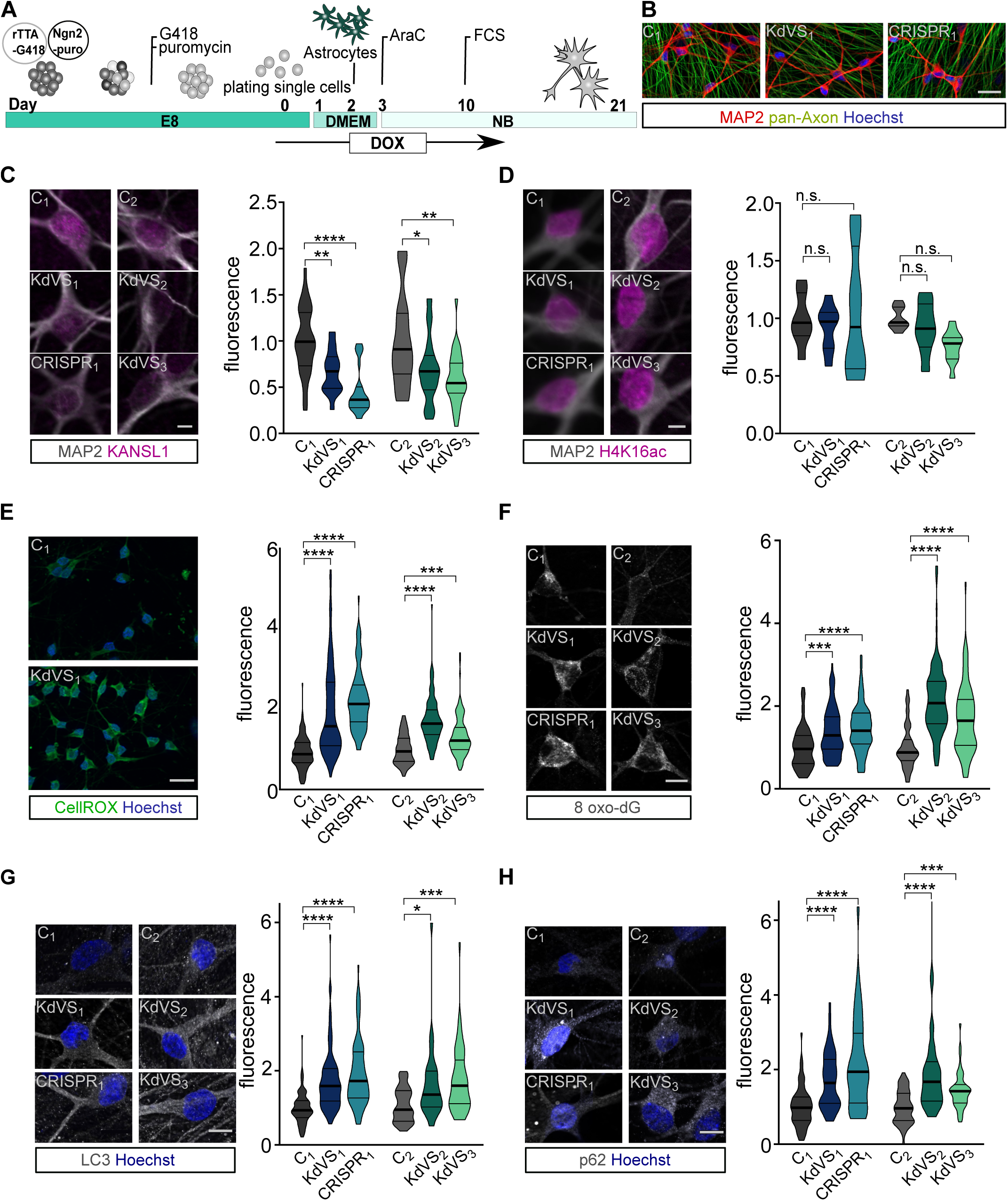
Autophagy phenotype in KdVS patient derived iNeurons. **A)** Schematic illustration of the protocol that was used for iPSC differentiation into neurons by means of doxycycline inducible Ngn2 expression. **B)** Representative images of iNeuron cultures at DIV21 stained for pan-axon and MAP2. Scale bar = 50 µm. **C)** Representative images of iNeurons at DIV21 stained for KANSL1 and MAP2. n = 33 for C_1_; n = 36 for KdVS_1_; n = 29 for CRISPR_1_; n = 26 for C_2_; n = 32 for KdVS_2_; n = 29 for KdVS_3_. Fluorescence quantification of nuclear KANSL1 signal was normalized to the respective control lines. Scale bar = 10 µm. **D)** Representative images of iNeurons at DIV21 stained for H4K16ac and MAP2. n = 7 for C_1_; n = 8 for KdVS_1_; n = 8 for CRISPR_1_; n = 8 for C_2_; n = 8 for KdVS_2_; n = 11 for KdVS_3_). Fluorescence quantification for H4K16ac immunostainings was normalized to the respective control lines. Scale bar = 10 µm. Significance was determined by means of one-way ANOVA and Sidak’s multiple comparison test. **E)** Representative images for CellROX staining for iNeurons derived of C_1_ and KdVS_1_ and fluorescence quantification for CellROX measurements at DIV21, normalized to the respective control lines. n = 125 for C_1_; n = 142 for KdVS_1_; n = 92 for CRISPR_1_; n = 78 for C_2_; n = 105 for KdVS_2_; n = 109 for KdVS_3_. Scale bar = 50 µm. **F)** Representative images for 8 oxo-dG stainings and fluorescence quantification for 8 oxo-dG stainings of iNeurons at DIV21 for all lines normalized to the respective control lines. n = 107 for C_1_; n = 102 for KdVS_1_; n = 80 for CRISPR_1_; n = 57 for C_2_; n = 105 for KdVS_2_; n = 86 for KdVS_3_. Scale bar = 10 µm. **G)** Representative images of LC3 immunostainings and fluorescence quantification relative to the respective control line lines. n = 90 for C_1_; n = 91 for KdVS_1_; n = 85 for CRISPR_1_; n = 39 for C_2_; n = 44 for KdVS_2_; n = 51 for KdVS_3_. Scale bar = 10 µm. **H)** Representative images of p62 stainings and fluorescence quantification relative to the respective control line lines. n=69 for C_1_; n = 60 for KdVS_1_; n = 85 for CRISPR_1_; n = 38 for C_2_; n = 42 for KdVS_2_; n = 47 for KdVS_3_. Scale bar = 10 µm. All data was obtained in at least 3 independent experiments. Statistically significant differences were tested through Kruskal-Wallis and Dunn’s multiple comparison test, if not mentioned differently. \**P*□<□0.05, \*\**P*□<□0.01, \*\*\**P*□<□0.005, \*\*\*\**P*□<□0.0001

Next, we sought to counteract the increased ROS levels by applying apocynin, a compound that prevents the formation of superoxide by suppressing NADPH oxidase. We were able to reduce ROS levels significantly in KdVS patient-derived iPSCs (Figure 2E). Importantly, this was accompanied with significantly lower p62 accumulation (Figure 2F), further suggesting that higher oxidative stress levels cause autophagosome accumulation in KANSL1-deficient iPSCs.

### ROS -induced increase in autophagosomes in KANSL1-deficient neurons

Dysregulated autophagy has been associated with neurodevelopmental disorders, including autism spectrum disorders^9,35–37^, and neurodegenerative diseases such as PD^38–41^ and amyotrophic lateral sclerosis (ALS)^42,43^, where familial cases are associated with mutations in *SOD1* ^44^. Autophagy is crucial to proper axon guidance, vesicular release, dendritic spine architecture, spine pruning, and synaptic plasticity^3,9,37,45,46^. Hence, we next examined whether the increase in autophagosomes observed in iPSCs is also present in KANSL1-deficient neurons, and whether a defective autophagy pathway affects specific aspects of neuronal function. To this end we differentiated iPSCs into a homogeneous population of excitatory cortical layer 2/3-like neurons (iNeurons) by forced expression of the transcription factor transgene Neurogenin-2 (*Ngn2)*^47,48^. For all experiments, iNeurons were co-cultured with freshly isolated rodent astrocytes to facilitate maturation (Figure 3A). All iPSC lines were able to differentiate into MAP2-positive neurons with comparable cell densities (Figure 3B, Supplementary Figure 5A). KANSL1 expression was lower in neurons derived from KANSL1-deficient iPSCs (Figure 3C) compared to controls. Similar to iPSCs, KANSL1-deficiency did not lead to a general reduction in H4K16ac in iNeurons (Figure 3D). Twenty-one days after the start of differentiation (days in vitro, DIV), both, control and KdVS iNeurons showed a fully developed neuronal morphology, measured by reconstruction and quantitative morphometry of MAP2-labeled iNeurons (Figure 3B, Supplementary Figure 4). KANSL1-deficiency did not result in any significant alteration in the neuronal somatodendritic morphology, including soma size, the number of primary dendrites, dendritic length and overall complexity (Supplementary Figure 4). Next, we measured ROS levels in DIV 21 iNeurons and found elevated ROS levels in KANSL1-deficient iNeurons (Figure 3E-F). To further corroborate our results, we immunostained iNeurons for 8-oxo-dG, an antibody targeting oxidized DNA, to assess levels of oxidative stress. KANSL1-deficient iNeurons all showed 50-100% increased 8-oxo-dG immunoreactivity (Figure 3F). This increase in ROS was also accompanied by increased levels of p62, LC3 and WIPI2 in KANSL1-deficient iNeurons at DIV21 (Figure 3G-H and Supplementary Figure 5B). Summarized, these results indicate that in KANSL1-deficient iNeurons ROS levels and the number of autophagosomes are increased, similar to our observations in KdVS patient-derived iPSCs.

**Figure 4.**
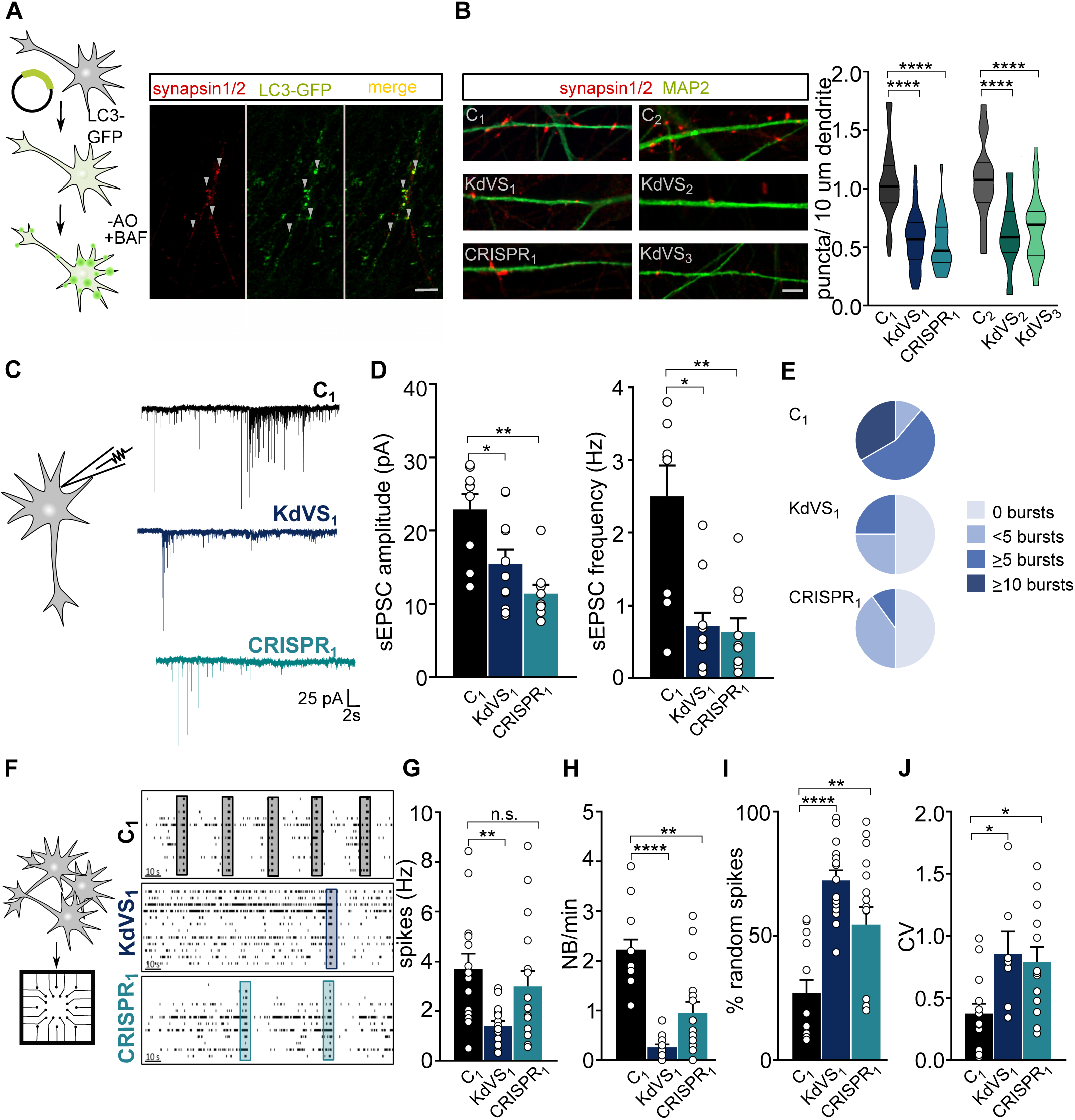
Synaptic phenotype in KdVS patient derived iNeurons. **A)** Schematic presentation of protocol to co-localize LC3 and synapsin. Image of a dendrite of control iNeuron at DIV21 after o/n incubation w/o B27 and treated with 200 nM Bafilomycin for 10 minutes before fixation. Scale bar = 10 µm. **B)** Representative images showing dendrites stained for MAP2 and synapsin 1/2 and synapsin puncta quantification at DIV21 for all lines. n = 60 for C_1_; n = 57 for KdVS_1_; n = 24 for CRISPR_1_; n = 15 for C_2_; n = 20 KdVS_2_; n = 34 for KdVS_3_. Scale bar = 20 µm. One-way ANOVA and Sidak’s multiple comparison test were used to test for statistically significant differences. **C)** Representative voltage clamp recordings at V_h_ = −60mV showing sEPSCs at DIV21. **D)** sEPSC amplitude and frequency quantification (n = 9 for C_1_; n = 11 for KdVS_1_; n = 10 for CRISPR_1_, obtained in two independent experiments). **E)** Percentages of cells for which 0 EPSC bursts, less than 5 EPSC bursts, 5 or more EPSC bursts, and 10 or more bursts were detected. **F)** Schematic representation for neuronal network measurements on MEAs (3 minutes of recording). Representative raster plots for C_1_, KdVS_1_ and CRISPR_1_ derived networks that were plated at similar high densities, measured at DIV 30. **G)** Quantification of the mean firing rate and **(H)** network burst rate, **(I)** percentage of random spikes, and **(J)** coefficient of variation (CV) calculated on the inter network burst interval. n = 15 for C_1_; n = 18 for KdVS_1_; n = 16 for CRISPR_1_. If not stated differently, data presented in this figure was obtained in at least three independent experiments and statistically significant differences were tested through Kruskal-Wallis and Dunn’s multiple comparison test. \**P*□<□0.05, \*\**P*□<□0.01, \*\*\**P*□<□0.005, \*\*\*\**P*□<□0.0001.

**Figure 5.**
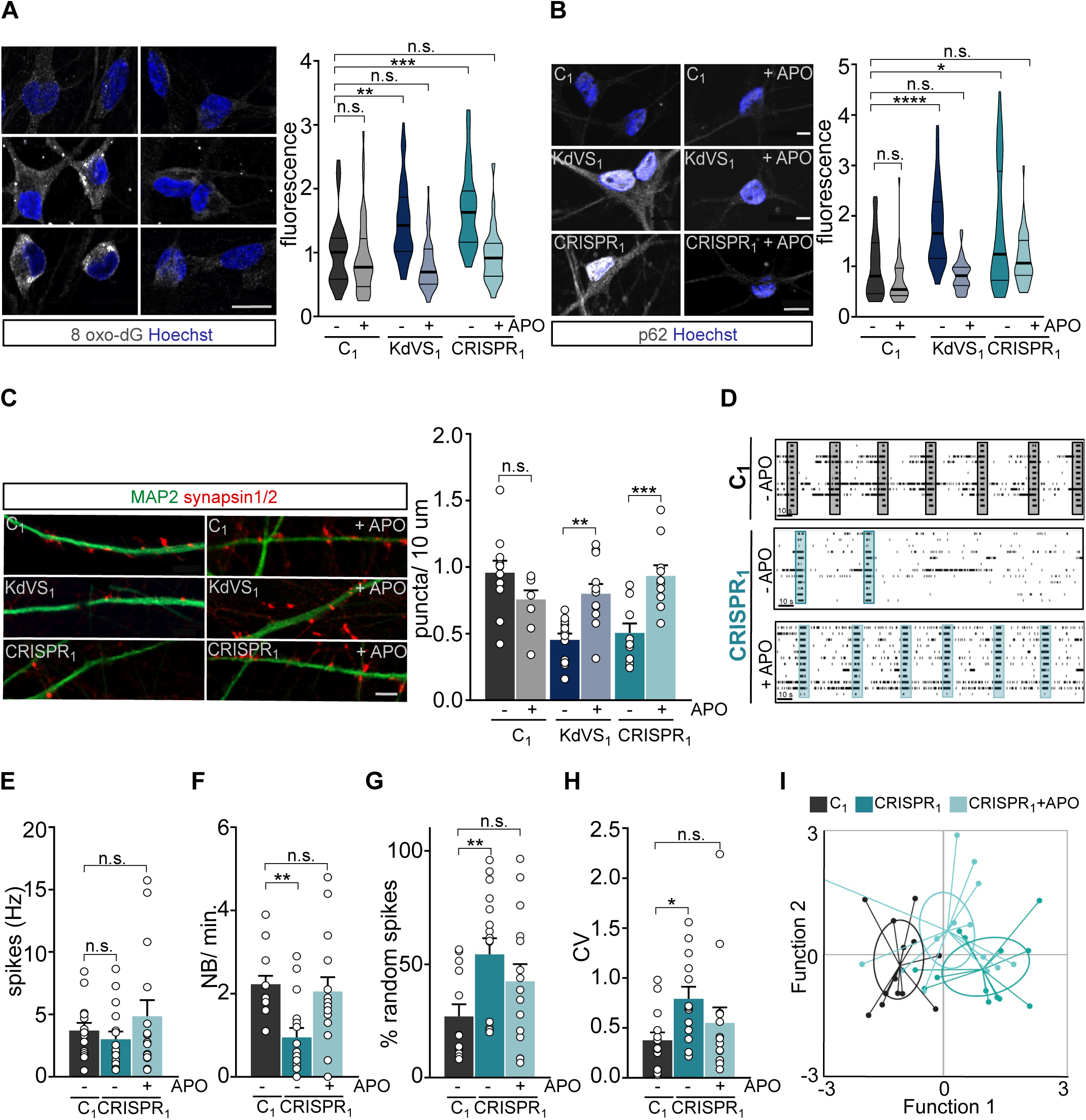
Apocynin treatment rescues synaptic phenotype. **A)** Representative images of 8 oxo-dG stainings and 8 oxo-dG quantification for untreated and APO treated iNeurons relative to untreated control cells at DIV21. n = 39 for C_1_; n = 46 for C_1_ +APO; n = 42 for KdVS_1_; n = 51 for KdVS_1_ +APO; n = 26 for CRISPR_1_; n = 25 for CRISPR_1_ +APO. Scale bar = 20 µm. **B)** Representative images of p62 stainings of iNeurons at DIV21 and fluorescence quantification in untreated and APO treated iNeurons relative to untreated control cells at DIV21. n = 43 for C_1_; n = 34 for C_1_+APO; n = 34 for KdVS_1_; n = 38 for KdVS1+APO; n = 40 for CRISPR_1_; n= 39 for CRISPR_1_+APO. Scale bar= 20µm. **C)** Representative images of dendrites stained for MAP2 and synapsin 1/2 for C_1_, KdVS_1_ and CRISPR_1_ at DIV21 either untreated or APO treated and synapsin quantification. n = 11 for C_1_; n = 9 for C_1_+APO; n = 12 for KdVS_1_; n = 12 for KdVS_1_+APO; n = 10 for CRISPR_1_; n = 10 for CRISPR_1_+APO (in two independent batches). Scale bar = 20µm. Two-way ANOVA was used to determine statistically significant changes. **D)** Representative raster plots for untreated C_1_ network and untreated and Apocynin treated CRISPR_1_ network at DIV30 (3 min. of recording). Quantification of **(E)** mean firing rate, **(F)** NB frequency, **(G)** percentage of random spikes, and **(H)** CV of inter-network burst interval. n = 15 for C_1_; n = 17 for C_1_ +APO; n = 16 for CRISPR_1_; n = 15 for CRISPR_1_ +APO. **I)** Canonical scores plot based on discriminant analyses for C_1_, CRISPR_1_ and CRISPR_1_ treated with APO. Discriminant functions are based on using the following network activity parameters: firing rate, network burst rate, network burst duration, percentage of spike outside network burst and coefficient of variability of the inter-network burst interval. Group envelopes (ellipses) are centered on the group centroids. All data presented in this figure were generated in at least 3 independent experiments and statistically significant differences were tested through Kruskal-Wallis and Dunn’s multiple comparison test, if not mentioned differently. \**P*□<□0.05, \*\**P*□<□0.01, \*\*\**P*□<□0.005, \*\*\*\**P*□<□0.0001.

### Decreased synaptic activity in KdVS patient derived iNeurons

Neurodevelopmental disorders have been associated with synaptic deficits. Also in the KdVS mouse model, KANSL1-deficiency has already been shown to result in impaired synaptic transmission^22^. Autophagosomes have been localized at the synapse, where mTOR-induced autophagy plays a role in synaptic pruning^9^. In addition, SOD1 and ROS have an essential role in synaptic function and memory^49–51^, but the link between ROS, autophagy and synaptic function remains unclear. Given our abovementioned observations we first investigated whether increased oxidative stress causes autophagosome formation at the synapse in iNeurons. To do so, we infected control iNeurons with a lentivirus to express LC3-GFP. At DIV21 we removed B27 from the medium in order to increase oxidative stress and treated the cells for 10 minutes with bafilomycin to prevent fusion of autophagosomes with lysosomes. Then we co-stained for synapsin 1/2. LC3 puncta co-localized with synapsin (Figure 4A), showing autophagosome formation at the synapse.

Next, we stained iNeurons for synapsin 1/2 at DIV21 and found significantly fewer synapsin puncta in KANSL1-deficient iNeurons compared to controls (Figure 4B), which indicates a reduced amount of putative functional synapses. To test whether there are indeed less functional synapses we performed whole-cell voltage-clamp recordings of spontaneous excitatory postsynaptic currents (sEPSCs) for three-week old C_1_-, KdVS_1_- and CRISPR_1_-derived iNeurons. KANSL1-deficient neurons (KdVS_1_ and CRISPR_1_) showed a clear reduction in amplitude and frequency of sEPSCs, compared to control C_1_ iNeurons (Figure 4C-D). Changes in frequency and amplitude suggest both, a pre- and postsynaptic deficit. Furthermore, we found that the KANSL1-deficient neurons received significantly less bursts of sEPSCs than controls (p<0.0001; Figure 4E), overall suggesting that KANSL1-deficient iNeurons received less excitatory input.

Dysfunction in neuronal network dynamics has been observed in the brain of patients with neurodevelopmental disorders^52–54^. In addition, neuronal network dysfunctions have been identified in model systems for several ID/ASD syndromes^55–57^. Our single-cell electrophysiological data prompted us to examine and compare the spontaneous electrophysiological population activity of control and KANSL1-deficient neuronal networks growing on micro-electrode arrays (MEAs). Through extracellular electrodes located at spatially-separated points across the cultures MEAs allowed us to monitor neuronal network activity non-invasively and repeatedly. As shown previously ^47^, electrical activity of control neuronal networks grown on MEAs organized into rhythmic, synchronous events (network burst), composed of numerous spikes occurring close in time and throughout all electrodes, four weeks after differentiation was induced (Figure 4F). This indicates that the iNeurons had self-organized into a synaptically-connected, spontaneously active network that matures over time. As expected from the reduced number of functional synapses, we observed significantly less frequent network bursts in networks composed of KdVS_1_-, as well as CRISPR_1_-derived iNeurons (Fig. 4F, H). The global level of spiking activity was significantly lower in networks of the KdVS patient-derived iNeurons (KdVS_1_), but not the CRISPR_1_ iNeurons (Figure 4G). However, we found that in both, the KdVS_1_ and CRISPR_1_ iNeurons, the percentage of random spikes was significantly increased (Figure 4I). Together with a decrease in network burst rate, this indicates that KANSL1-lacking networks organized differently compared to controls, in line with a more immature state in which activity mainly occurs outside network bursts^47,55^ (Figure 4I). This was further illustrated by a more irregular network-bursting pattern, quantified as a larger coefficient of variation (CV) (Figure 4J) of the inter-network burst interval for KdVS_1_- and CRISPR_1_-derived networks.

### Apocynin treatment rescues synaptic phenotype in KdVS derived neurons

To investigate whether increased oxidative stress is causal to the autophagosome formation and subsequent reduced synaptic activity in KdVS patient-derived iNeurons we treated iNeurons with 100 µM apocynin. First, we confirmed that prolonged apocynin treatment reduced ROS levels and autophagosome accumulation in KANSL1-deficient iNeurons of KdVS_1_ and CRISPR_1_, through 8 oxo-dG and p62 immunostainings, respectively (Fig. 5 A-B). In control line C_1_ apocynin treatment did not change 8 oxo-dG and p62 levels significantly. Having reduced ROS in KdVS iNeurons with apocynin we next examined the effect on synapse density. Prolonged apocynin treatment (for 14 days) increased synapsin puncta significantly in KdVS iNeurons, to a level that was comparable to control iNeurons (Figure 5C). Of note, apocynin treatment had little effect on control iNeurons, suggesting that ROS levels in normal circumstances are low in these cells. In order to show that the increase in synapsin puncta correlates with an increase in synaptic activity we measured neuronal network activity for apocynin-treated iNeurons. As described previously, neuronal network bursts occurred significantly less often in KANSL1-deficient networks. Reducing oxidative stress by means of apocynin treatment significantly increased the number of network bursts for CRISPR_1_ to control level (Fig. 5 D, F). This was further accompanied with unaltered mean firing rate, a reduction in the percentage of random spikes and a reduction in the CV of the inter-network burst interval, indicating that the network pattern became more regular and mature, similar to control networks (Figure 5 E-H). This is also illustrated by canonical scores plots based on discriminant analyses for C_1_, CRISPR_1_ and CRISPR_1_ treated with apocynin (Fig. 5I) and reclassification of group membership (Supplementary Figure 6).

**Figure 6.**
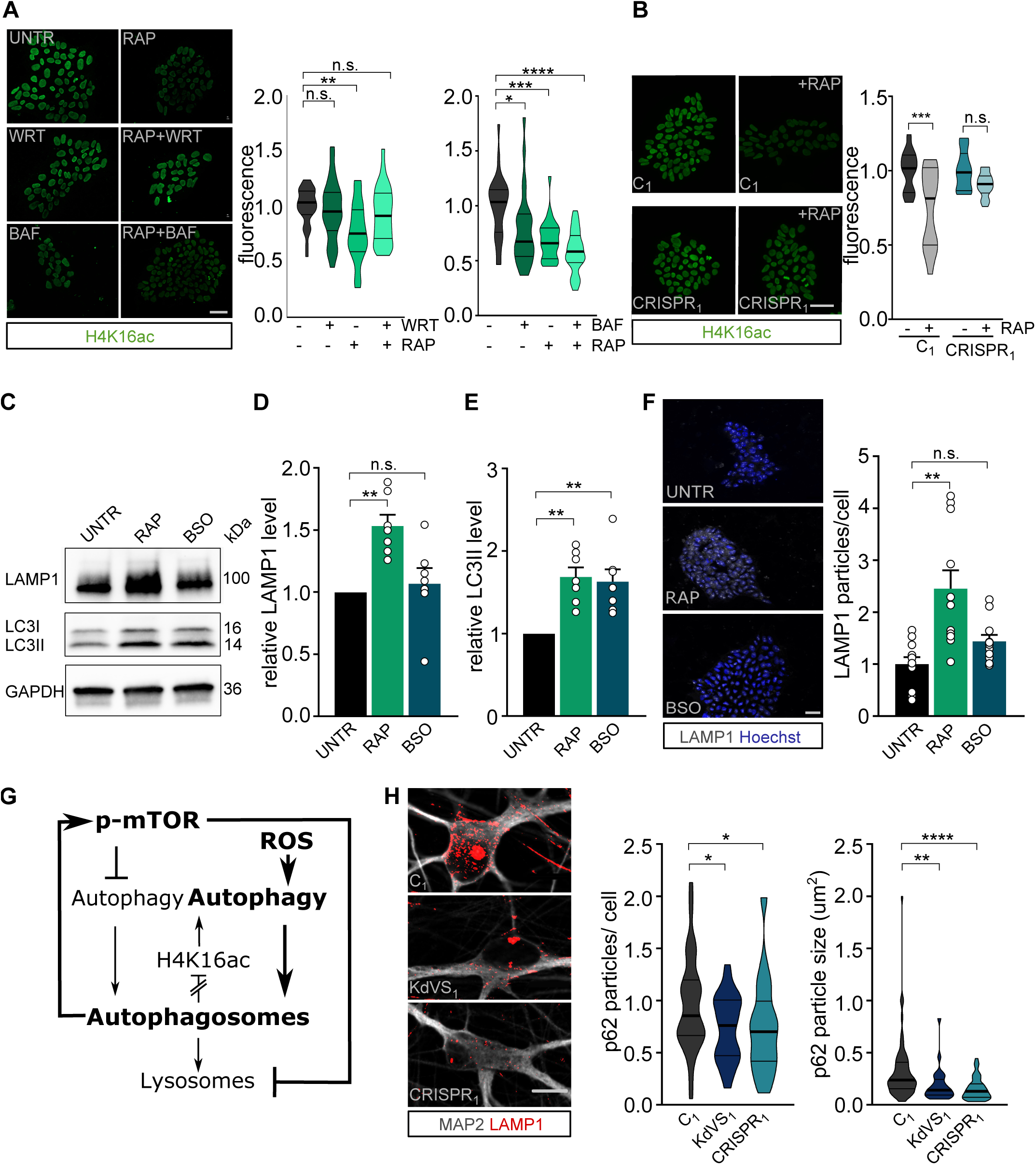
Feedback loop activation and reduced lysosomal activity. **A)** Representative images of iPSCs treated with Rapamycin (RAP) and/or Wortmannin (WRT) or Bafilomycin (BAF) stained for H4K16ac. Fluorescence was always normalized to untreated control samples. n = 29 for RAP/WRT treatments; n = 25 for RAP/BAF treatments. Two-way ANOVA and Tukey’s multiple comparison test were used to test for statistically significant differences. Scale bar = 50 µm. **B)** Representative images of iPSC colonies of C_1_ and CRISPR_1_ untreated or treated with RAP (200 µM) for 10 minutes. Two hours after the treatment cells were fixed and stained for H4K16ac. n = 12 for C_1_; n = 15 for C_1_ +RAP; n = 10 for CRISPR_1_; n = 14 for CRISPR_1_ +RAP. Two-way ANOVA and Sidak’s multiple comparison test were used to determine statistically significant reduction. Scale bar = 50 µm. **C)** Representative Western blot for LAMP1 and LC3 after autophagy induction for 10 minutes by either RAP (200 µM) or BSO (200 µM) in control iPSCs. **D)** LAMP1 protein level quantification and **(E)** LC3II quantification relative to untreated control. n = 7 for all conditions. Kruskal Wallis and Dunn’s multiple comparison test were used to test for statistically significant differences. **F)** Representative images of LAMP1 stainings in control iPSCs treated with RAP (200 µM) or BSO (200 µM) for 10 minutes before fixation and particle analysis for LAMP1. n = 11 for all conditions. Scale bar = 50 µm. One-way ANOVA and Dunnett’s multiple comparison test were used to determine statistically significant differences. **G)** Schematic representation of autophagy showing the 2 different autophagy inducing pathways discussed (mTOR and ROS). NSL complex mediated feedback-loop is induced by autophagosome formation. At the same time mTOR phosphorylation is increased, which subsequently reduces lysosomal activity.LC3II protein level quantification. **H)** Representative images of three-week old neurons from C_1_, KdVS_1_ and CRISPR_1_ stained for MAP2 and LAMP1 and LAMP1 particle quantification. n = 43 for C_1_; n = 29 for KdVS_1_; n = 36 for CRISPR_1_. Results were normalized to control. Scale bar = 10 µm. One-way ANOVA and Dunnett’s multiple comparison test were used to determine significant differences for the number of particles. Differences in particle size were tested through Kruskal Wallis and Dunn’s multiple comparison test. Data presented in this figure was obtained in at least two independent experiments. \**P*□<□0.05, \*\**P*□<□0.01, \*\*\**P*□<□0.005, \*\*\*\**P*□<□0.0001.

### Impaired feedback-loop and decreased lysosomal activity in KdVS cells

Having established that KANSL1-deficiency results in increased oxidative stress causing elevated autophagosome formation and reduced synaptic activity, we wondered what causes the autophagosome accumulation in KANSL1-deficient cells. Increased autophagic flux is known to induce a regulatory feedback-loop in which induction of autophagy is coupled to a reduction in H4K16ac through downregulation of hMOF in MEF cells^15^. However, we did not find a reduction in H4K16ac in KANSL1-deficient cells at the basal level. This led to the question, whether an impaired feedback-loop activation via the NSL complex is underlying the observed autophagosome accumulation.

To answer this question, we first examined at which step during autophagy signaling the negative feedback-loop is activated. After activation there are three main stages of autophagy: (i) the formation of initiation complexes, (ii) autophagosome formation and (iii) fusion with lysosomes. We pharmacologically inhibited the different stages within the autophagy pathway to identify which stage triggers the feedback-loop activation in control line C_2_. Blocking autophagosome formation with wortmannin, a PI3-kinase inhibitor, did not result in reduced H4K16ac after rapamycin treatment (Figure 6A). However, blocking lysosome fusion with bafilomycin resulted in activation of the negative feedback-loop (Figure 6A), indicated by reduced levels of H4K16ac. This suggests that the formation of autophagosomes is the signal for the feedback-loop activation. Notably, the reduction in H4K16ac was independent of the autophagy induction protocol, since a similar decrease in H4K16ac was observed when iNeurons were stimulated with buthionine sulfoximine (BSO) in order to activate ROS-mediated autophagy^58^ (Supplementary Figure 7A). In contrast to wild type cells, autophagosome accumulation in KANSL1-deficient cells did not result in reduced H4K16ac, pointing towards changes in NSL complex-mediated feedback-loop activation (Figure 6B, Supplementary Figure 7A). Although we did not observe reduced H4K16ac, we found increased mTOR phosphorylation (Figure 2A, F) in KANSL1-deficient iPSCs, which might be part of a negative feedback-loop that is activated independently from KANSL1 and the NSL complex to prevent prolonged autophagy activation. These findings point towards a tightly controlled, balanced interaction between the different autophagy regulating mechanisms. Autophagosome formation induces independent signaling cascades to negatively regulate autophagy through, on the one hand, increased mTOR activity, and, on the other hand, reduced H4K16ac levels to repress ATG gene expression. In KANSL1-deficient cells autophagy regulation is out of balance; H4K16ac is not reduced, enabling prolonged ROS activated autophagy.

**Figure 7.**
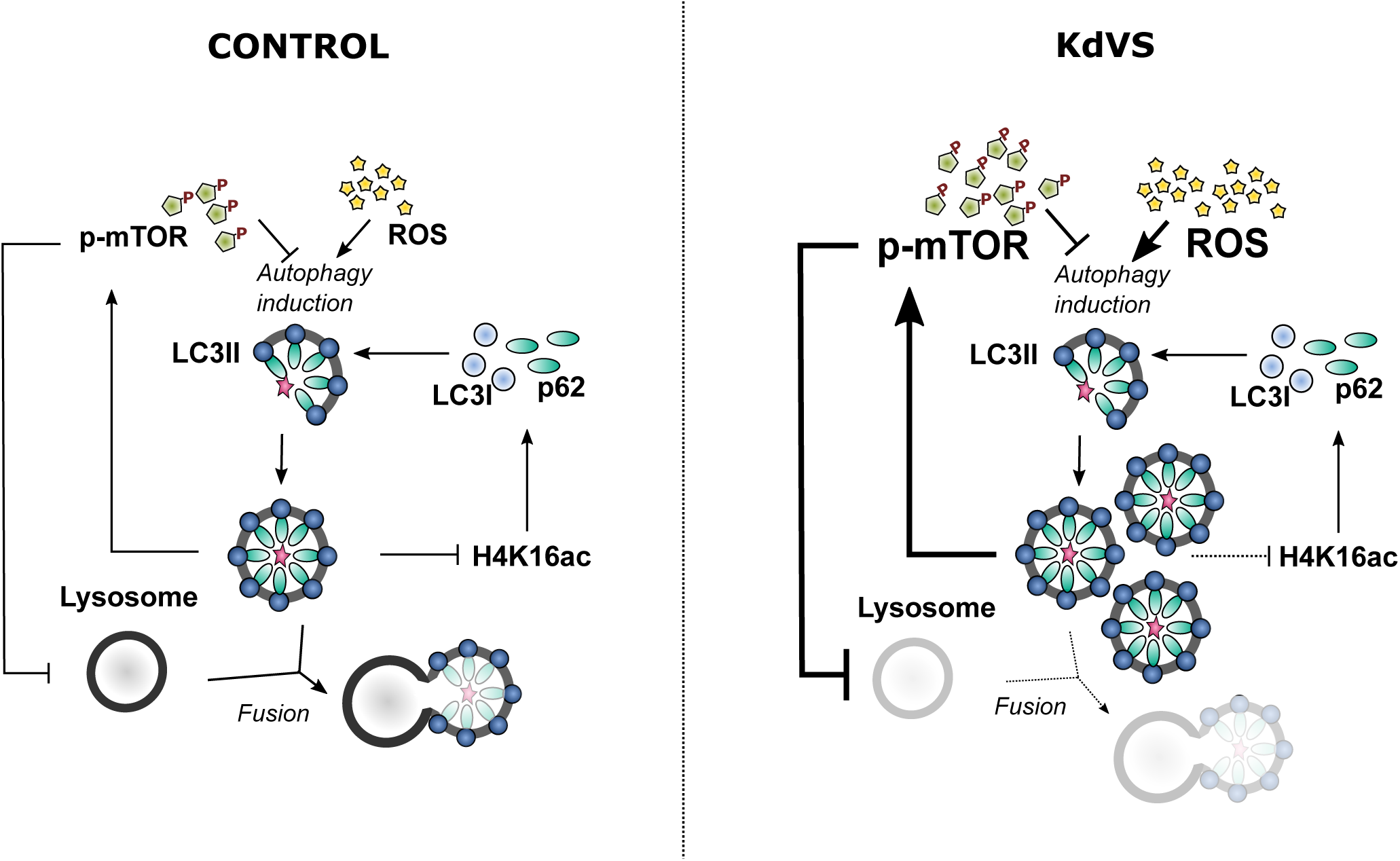
Autophagy Regulation in control and KdVS cells. Schematic representation of autophagy regulating mechanisms. In control cells both, dephosphorylation of mTOR and increased ROS levels, induce autophagosome formation. Upon completion, autophagosomes induce negative feedback-loops. ATG gene expression is reduced through decreased H4K16ac levels, on the one hand. On the other hand, mTOR phosphorylation increases upon autophagosome accumulation to inhibit mTOR regulated autophagy, and subsequently reduces lysosomes. In KdVS cells prolonged oxidative stress induces autophagy, primarily mTOR independent. While H4K16ac is not reduced, enabling expression of ATG genes and continuous autophagosome formation, the accumulation of autophagosomes induces a reduction in mTOR activity, reducing lysosomal activity, and preventing autophagosomal clean-up. Aberrant feedback-loop activation in KdVS cells therefore causes an imbalance in oxidative stress versus mTOR activated autophagy reinforcing elevated autophagosome accumulation.

Since active mTOR signaling is known to decrease lysosomal activity^59–61^, we tested whether lysosomal activity might be affected by the imbalanced autophagy regulation in KANSL1-deficient cells. First, we quantified LAMP1, a lysosomal marker, in rapamycin treated iPSCs by means of immunocytochemistry and Western blot. We confirmed that mTOR-dependent autophagy activation increases autophagosome formation (Figure 6C, E) as well as lysosomal activity (Figure 6C-D, F) in control cells. BSO treatment, to activate ROS-induced autophagy, increased LC3II levels comparable to rapamycin treatment (Figure 6C, E), whereas LAMP1 levels remained low (Figure 6C-D, F). We found increased autophagosome formation in KANSL1-deficient cells to be initiated by increased ROS, which points towards less efficient lysosome activation. Indeed, quantifying LAMP1 particles in KANSL1-deficient iNeurons indicated significantly lower lysosomal activity in these cells when compared to control iNeurons (Figure 6H). Additionally, decreased ROS levels through apocynin treatment showed to reduce the p-mTOR/ mTOR ratio and also increased LAMP1 levels in KANSL1 deficient iPSCs (Supplementary Figure 7B-D). Together, our data thus suggest that increased ROS levels activate autophagy. Subsequently, autophagosome formation induces mTOR phosphorylation leading to reduced lysosomal activity which reinforces autophagosome accumulation in KANSL1-deficient iNeurons (Figure 6G, 7).

## Discussion

In this study we used a human *in vitro* model for KdVS to examine the role for KANSL1 in autophagy regulation and to gain insight into how deregulated autophagy affects neuronal function. We found that KANSL1-deficiency leads to increased oxidative stress and autophagosome formation in iPSCs and iNeurons (Figure 7). In neurons increased ROS activated autophagy showed to reduce neuronal synaptic connectivity and activity, revealed on single cell, as well as on network level. The observed neuronal phenotype could be rescued by treatment with apocynin, an antioxidant that reduced oxidative stress and autophagosome accumulation.

### Imbalanced mTOR- and ROS-mediated autophagy in KdVS cells

Heterozygous loss of *KANSL1* lead to imbalanced autophagy regulation in KANSL1-deficient iPSCs and neurons. On the one hand, we found increased ROS-mediated autophagy, and, on the other hand, we found reduced mTOR-mediated autophagy. Previous studies in neurons have mostly associated deregulated autophagy to increased or decreased activity of mTOR-regulated autophagy^9,36,62^. Here, we show that the interplay between both autophagy pathways is essential for balanced autophagy regulation. Little is known about the interaction between the different autophagy pathways and how exactly the feedback-loop is controlled. It has been reported that upon autophagy induction, H4K16ac is downregulated in order to reduce ATG gene expression and prevent prolonged autophagy^15^. Here, we provide more insight in this feedback-loop activation by showing that in control iPSCs acute autophagosome formation induces H4K16ac reduction within a couple of hours. In contrast, acute activation of autophagy in KANSL1-deficient cells did not result in a significant H4K16ac reduction. This is in line with our observation that ROS-mediated accumulation of autophagosomes in KANSL1-deficient cells (iPSCs and iNeurons) did not affect H4K16ac levels when compared to control cells. According to what is known about the role of H4K16ac in autophagy regulation, and especially in negative feedback-loop activation, this should eventually lead to cell death^15^. However, we cultured KANSL1-deficient iNeurons for three weeks and longer without observing increased cell death. This indicates that there might be an additional feedback mechanism active that is inhibiting aspects of prolonged autophagy that would otherwise lead to cell death, independent from the NSL complex mediated feedback^63^. We observed increased mTOR activity in KANSL1-deficient iPSCs, which we interpreted as possible feedback reaction to counteract hyperactive ROS-induced autophagy. Upon starvation induced autophagy, mTOR is first deactivated, but reactivated shortly after autophagic degradation, even during prolonged starvation. mTOR activation is known to inhibit the expression of lysosomal genes by phosphorylating the transcription factor TFEB and preventing its’ nuclear translocation^60,64,65^. In a model for Gaucher’s disease, mTOR hyperactivity was shown to cause lysosomal dysfunction in neurons^66^. Similarly, we found increased levels of active mTOR (p-mTOR) and reduced levels of the lysosomal marker LAMP1 in KANSL1-deficient iNeurons. This indicates that hyperactive ROS-mediated autophagosome formation initiates a feedback-loop that increases mTOR activity which subsequently blocks lysosomal gene expression in KANSL1-deficient cells. Interestingly, valproic acid treatment, a histone deacetylase inhibitor, not only prevents H4K16ac downregulation after autophagy induction, but also increases autolysosome formation and causes cell death^15^. Combining these findings with our observation that we did not observe increased cell death; we suggest that activation of mTOR in KANSL1-deficient cells is required to avoid prolonged lysosomal degradation and subsequent cell death. While our results show that rapamycin treatment can efficiently induce autophagosome clearance within a few hours, the interplay between oxidative stress- and mTOR-mediated autophagy indicate that long-term rapamycin treatment as a therapeutic strategy is questionable. Continuous over-activation of even two autophagy activating pathways in combination with lacking negative feedback, would probably lead to cell death in KANSL1 deficient cells. In contrast, we find that preventing ROS production with apocynin was sufficient to rescue the autophagy phenotypes. More importantly, apocynin was sufficient to rescue normal neuronal functioning, suggesting that increased ROS is causal to the autophagy phenotypes and at the core of KdVS pathophysiology.

Loss of function mutations in *KANSL1* have unequivocally been associated with KdVS. More recently, two single nucleotide polymorphism (SNP) in *KANSL1* (Ile1085Thr and Ser718Pro) have been identified as risk factors for PD in a genome-wide association study (GWAS)^67^. This is of interest since it is well established that both, oxidative stress and autophagy (and mitophagy) deficits, are strongly associated with PD and other neurodegenerative disorders^68–70^. Furthermore, it is widely accepted that ROS activate autophagy to restore cellular homeostasis as a cytoprotective feedback mechanism. Evidence from ageing and neurodegenerative models indicate impaired neuronal function and ultimate neuronal cell death when this regulatory link is disturbed by either reduced autophagic flux, or elevated ROS formation exceeding capacity for cytoprotection^71^. Our results provide additional evidence for the crucial link between ROS and autophagy for regulation of neuronal cell homeostasis and function, not only during neurodegeneration but also in the context of a neurodevelopmental disorder. Since the SNPs in *KANSL1* identified in PD GWAS are unlikely to lead to loss-of-function it will be of particular interest to investigate the functional consequences of these SNPs in autophagy.

### ROS-induced autophagy regulates synaptic function

Whereas autophagy has been extensively studied in the context of neurodegenerative disorders, its’ role during neuronal development has remained understudied. Especially how mTOR-independent autophagy affects synaptic development and function is largely unknown. This is of relevance since it is known that levels of ROS, or more specifically superoxide, at the synapse plays a crucial role in synaptic plasticity^72^. On the one hand, local increase in superoxide levels upon neuronal stimulation showed to be essential to induce hippocampal long-term potentiation (LTP)^73–75^. On the other hand, increased levels of superoxide contribute to age-related impairments in hippocampal LTP and memory^76^. Our single-cell electrophysiology data showed reduced synaptic input, i.e. reduced sEPSC frequency and less frequent sEPSC bursts. In addition, we observed a significantly reduced sEPSC amplitude pointing towards a reduction in functional AMPA receptors at the synapse. This indicates that chronic, ROS-induced autophagy might affect receptor distribution at the post-synaptic site. It is already known that chemical induction of long-term depression (LTD) induces NMDAR-dependent autophagy through mTOR inhibition, resulting in AMPAR break down^11^. These results further support the evidence that ROS-mediated autophagy is contributing to the regulation of synaptic plasticity. ROS-mediated changes during synaptic plasticity might occur through the activation of autophagy to induce synapse-specific break-down of synaptic proteins. More detailed investigation is, however, needed to understand how synaptic input- and target-specificity (e.g. AMPAR subunits) is achieved during autophagy at synapses in light of synaptic plasticity.

Taken together our findings establish a previously unrecognized link between H4K16ac, ROS-mediated autophagy, synaptic dysfunction and KdVS. Future research should further examine the role of autophagy in neurodevelopment, focusing on the interplay between the different autophagy induction pathways and its fin-tuned regulation through epigenetic mechanisms.

## Material and Methods

### Patient information and iPSC line generation

In this study we used two control and three KdVS patient derived iPSC lines (Figure 1). All patients that were included in this study present the full spectrum of KdVS associated symptoms. KdVS_1_^19^ and KdVS_2_^17,77^ originate from two individual, female KdVS patients with mutations in KANSL1, while the iPSC line KdVs_3_ originates from a female patient with a 17q microdeletion^19,78^. We used one independent female control lines (C_2_). Next to this control line we also used a parent control line for patient line KdVS_1_ (C_1_). C_1_ was also used for CRIPSR/Cas9 genomic engineering in order to create a *KANSL1* mutated line with the same genetic background (CRISPR_1_). A 1 bp insertion in Exon 2 resulted in a premature stop codon and heterozygous loss of *KANSL1*, highly similar to the patient lines KdVS_1_ and KdVS_2_.

All iPSCs used in this study were obtained from reprogrammed fibroblasts. KdVs_1_ and C_1_ were generated by making use of the Simplicon™ reprogramming kit (Millipore). Overexpression of the four factors Oct4, Klf4, Sox2, and Glis1 was introduced by a non-integrative, non-viral one step transfection. KdVS_2_ was reprogrammed by lentiviral mediated overexpression of Oct4, Sox2, Klf4, and cMyc. For KdVS_3_ episomal reprogramming was performed with the same reprogramming factors. C_2_ was generated from fibroblasts of a 36-year-old female control obtained from the Riken BRC – Cell engineering division (HPS0076:409B2). iPSC clones used in this study were validated through a battery of quality control tests including morphological assessment and karyotyping to confirm genetic integrity. All clones expressed the stem cell markers OCT4, SOX2 NANOG, SSEA-4, and TRA1-81 (see supplementary Figure 1 A).

### Genome editing by CRISPR/Cas9

We performed CRISPR/Cas9 genome editing on C_1_ following the protocol from Ran et al. (2013). First sgRNA targeting exon 2 of *KANSL1* was cloned into the targeting vector (pX459v2). Successful cloning was validated by PCR. Single iPSCs were then nucleofected with the pX459v2 plasmid coding for the Cas9 protein and the sgRNA using P3 primary Cell 4D nucleofector kit (Lonza, V4XP-302). After nucleofection cells were plated on a 6-well plate in E8 flex medium supplemented with RevitaCell. 24 Hours after nucleofection medium was refreshed and supplemented with puromycin (1 µg/ml), RevitaCell was removed. After o/n incubation medium was refreshed with E8 flex (Thermo Fisher Scientific) to remove dead cells and stop the selection. The culture was maintained until colonies were large enough to be picked. By means of Sanger sequencing we checked for heterozygous loss-of-function mutations. Positive colonies were re-plated as single cells to ensure clonal expansion of iPSCs positive for the selected mutation. KANSL1 sgRNA oligos: 5’-CACCG**GAGCCCGTTTTCCCCCATTG**-3’; 3’-C**CTCGGGCAAAAGGGGGTAAC**CAAA-5’

### Generation of rtTA/Ngn2-positive iPSCs

For the generation of rtTA/ Ngn2-positive iPSCs, lentiviral vectors were used to stably integrate the transgenes into the genome of the iPSCs. The vector used for the rtTA lentivirus is pLVX-EF1α-(Tet-On-Advanced)-IRES-G418(R); it encodes a Tet-On Advanced transactivator under control of a constitutive EF1α promoter and confers resistance to the antibiotic G418. The lentiviral vector for murine neurogenin-2 (*Ngn2*) was pLVX-(TRE-tight)- (MOUSE)Ngn2-PGK-Puromycin(R), encoding for *Ngn2* under control of a Tet-controlled promoter and the puromycin resistance gene under control of a constitutive PGK promoter. Both vectors were packaged into lentiviral particles using the packaging vectors psPAX2 lentiviral packaging vector (Addgene #12260) and pMD2.G lentiviral packaging vector (Addgene #12259). We infected our iPSC lines with both lentiviruses in order to ensure Ngn2 expression when the medium was supplemented with doxycycline. To select for iPSCs that were transduced with both vectors we started G418 (0.5 µg/ml) and puromycin (0.1 ug/ml, Invivogen) selection 48 hours after infection. The antibiotics concentration was doubled at day two and three of the selection process. iPSCs surviving the selection process were cultured at general iPSC culture conditions (see below).

### iPSC Culture and drug treatment

iPSCs were always cultured on Matrigel (Corning, #356237) in E8 flex (Thermo Fisher Scientific) supplemented with primocin (0.1 µg/ml, Invivogen) and low puromycin and G418 concentrations (0.5 µg/ml) at 37°C/5% CO^2^. Medium was refreshed every 2-3 days and cells were passaged 1-2 times per week using an enzyme-free reagent (ReLeSR, Stem Cell Technologies). For autophagy induction cells were treated with 200 µM Rapamycin (ChemCruz) for 10 minutes, before medium was refreshed. To block autophagosome formation or lysosome fusion cells were treated with 200 nM Wortmannin (Invivogen) or Bafilomycin (Millipore), respectively prior Rapamycin treatment. If not mentioned differently, cells were lysed/ fixated after 2 hours of incubation. For Apocynin (Santa Cruz Biotechnology) rescue experiments iPSCs were plated as single cells. The day after plating the cells were treated for 24 hours with 100 µM Apocynin before cells were fixed. To stimulate ROS production cells were treated for 10 minutes with 100 µM BSO (Sigma). After medium was refreshed, cells were incubated for 2 hours before lysate preparation or fixation.

### Neuronal differentiation

iPSCs were directly derived into upper-layer, excitatory cortical neurons by doxycycline induced overexpression of *Ngn2* according to an already published protocol^47^. To start neuronal differentiation Accutase (Sigma) was applied to generate single cells. iPSCs were then plated onto MEAs (600 cells/mm^2^) or glass, nitric-acid treated coverslips (100 cells/mm^2^) in E8 basal medium (Thermo Fisher Scientific) supplemented with primocin (0.1 µg/ml, Invivogen), RevitaCell (Thermo Fisher Scientific), and doxycycline (4µg/ml). MEA plates, as well as coverslips, were pre-coated with Poly-L-Ornithine (50 µg/mL, Sigma) for 3 hours at 37 °C and 20 µg/mL human laminin (L521, Biolamina) overnight (o/n) at 4 °C. The day after plating medium was changed to DMEM-F12 (Thermo Fisher Scientific) supplemented with N2 (Thermo Fisher Scientific), NT3 (Promokine), BDNF (Promokine), NEAAS (Sigma), doxycycline (4µg/ml), and primocin (0.1 µg/ml). To support neuronal maturation, freshly prepared rat astrocytes were added to the culture in a 1:1 ratio two days after plating. At DIV 3 the medium was changed to Neurobasal medium (Thermo Fisher Scientific) supplemented with B-27 (Thermo Fisher), glutaMAX (Thermo Fisher), primocin (0.1 µg/ml), NT3, BDNF, and doxycycline (4 µg/ml). Cytosine β-D-arabinofuranoside (Ara-C) (2 µM; Sigma) was added once to remove proliferating cells from the culture. From DIV 6 onwards half of the medium was refreshed three times a week. Addition of doxycycline was stopped after two weeks to reduce stress. The medium was additionally supplemented with 2,5% FBS (Sigma) to support astrocyte viability from DIV10 onwards. Neuronal cultures were kept through the whole differentiation process at 37°C/ 5%CO_2_. For rescue experiments cells were treated with 100 µM Apocynin every other day when medium was not refreshed from DIV6 onwards.

### Immunocytochemistry

Cells were fixed with 4% paraformaldehyde/ 4% sucrose (v/v) and permeabilized with 0.2% triton in PBS for 10 minutes. Aspecific binding sites were blocked by incubation in blocking buffer (PBS, 5% normal goat serum, 5% normal horse serum, 5% normal donkey serum, 1% bovine serum albumin, 1% Glycine, 0.2% Triton) for 1 hour at room temperature (RT). Primary antibodies were diluted in blocking buffer for o/n incubation at 4°C. Secondary antibodies, conjugated to Alexa-fluorochromes, were also diluted in blocking buffer and applied for 1 hour at RT. Hoechst was used to stain the nucleus before cells were mounted using DAKO (DAKO) fluorescent mounting medium and stored at 4°C. Used primary antibodies were: mouse anti-MAP2 (1:1000; Sigma M4403); guinea pig anti-MAP2 (1:1000; Synaptic Systems 188004); guinea pig anti-synapsin ½ (1:1000; Synaptic Systems 106004); rabbit anti-PSD95 (1:50; Cell Signaling D27E11); mouse anti-pan axon (1:1000; Covance SMI-312R); rabbit anti-p62 (1:500; Sigma p0067); mouse anti-LC3 (1:500; NanoTools 0231-100/LC3-5F10); rabbit anti-LAMP1 (1:200; L1418-200ul); mouse anti 8-oxo-dG (1:100; R&D Systems 4354-MC-050). Secondary antibodies that were used are: goat anti-guinea pig Alexa Fluor 647 (1:2000, Invitrogen A-21450); goat anti-rabbit Alexa Fluor 488 (1:2000, Invitrogen A-11034); goat anti-rabbit Alexa Fluor 568 (1:2000, Invitrogen A-11036); goat anti-mouse Alexa Fluor 488 (1:2000, Invitrogen A-11029); goat anti-mouse Alexa Fluor 568 (1:2000, Invitrogen A-11031). Cells were imaged with the Zeiss Axio Imager 2 equipped with apotome. All conditions within a batch were acquired with the same settings in order to compare signal intensities between different experimental conditions. Signals were quantified using FIJI software. The number of synaptic puncta was quantified per individual cell via manual counting and divided by the dendritic length of the neuron. For p62 and LC3 puncta quantification the Particle Analyzer tool was used.

### Oxidative stress quantification

To quantify oxidative stress/ ROS we used either 8 oxo-dG stainings or CellROX® assays (Thermo Fisher; Green Reagent). 8 oxo-dG (8-oxo-2’-deoxyguanosine) is an oxidized derivative of deoxyguanosine, one of the major products of DNA oxidation. Quantifying 8 oxo-dG can therefore be used to measure oxidative stress levels. CellROX® Green Reagent is fluorogenic, cell-permeant dye which is weakly fluorescent in a reduced state and exhibits bright green photostable fluorescence upon oxidation by ROS. Quantifying fluorescence therefore is an indication for oxidative stress. For both assays, cells were prepared according to previously mentioned protocols. For 8 oxo-dG we performed ICC on PFA fixed cells with a specific antibody (R&D System) to quantify DNA oxidation. For CellROX measurements we added the probe (5µM) to living cells and incubated for 30 minutes before fixation. After fixation the cells were mounted with DAKO and imaged after o/n incubation within 24 hours with the Zeiss Axio Imager 2 w/o apotome. Fluorescence was quantified using FIJI image software.

### Seahorse Mito Stress Test

Oxygen consumption rates (OCR) were measured using the Seahorse XFe96 Extracellular Flux analyzer (Seahorse Bioscience). iPSCs were seeded at a concentration of 10,000 per well in E8 basal medium supplemented with primocin, 10 µg/mL RevitaCell, and allowed to adhere at 37°C and 5% CO2. The day after plating RevitaCell was removed from the medium. One hour before measurement, culture medium was removed and replaced by Agilent Seahorse XF Base Medium (Agilent) supplemented with 10 mM glucose (Sigma), 1 mM sodium pyruvate (Gibco), and 200 mM L-glutamine (Life Sciences) and incubated at 37°C without CO_2_. Basal oxygen consumption was measured six times followed by three measurements after each addition of 1 µM of oligomycin A (Sigma), 2 µM carbonyl cyanide 4-(trifluoromethoxy) phenylhydrazone FCCP (Sigma), and 0.5 µM of rotenone (Sigma) and 0.5 µM of antimycin A (Sigma), respectively. One measuring cycle consisted of 3 minutes of mixing, 3 minutes of waiting and 3 minutes of measuring. The OCR was normalized to citrate synthase activity, to correct for the mitochondrial content of the samples^79^. The citrate synthase activity was measured according to the protocol described by Srere et al.^80^, modified for Seahorse 96 wells plates, as previously reported ^81^. In short, after completion of OCR measurements the Seahorse medium was replaced by 0.33% Triton X-100, 10 mM Tris-HCl (pH 7.6), after which the plates were stored at −80°C. Before measurements, the plates were thawed and 3 mM acetyl-CoA, 1 mM DTNB, and 10% Triton X-100 was added. The background conversion of DTNB was measured spectrophotometrically at 412 nm and 37 °C for 10 minutes at 1-minute intervals, using a Tecan Spark spectrophotometer. Subsequently, the reaction was started by adding 10 mM of the citrate synthase substrate oxaloacetate, after which the ΔA412 nm was measured again for 10 minutes at 1-minute intervals. The citrate synthase activity was calculated from the rate of DTNB conversion in the presence of oxaloacetate, subtracted by the background DTNB conversion rate, using an extinction coefficient of 0.0136 µmol-1. cm-1.

### Neuronal morphology analysis

To examine dendritic morphology of neurons, cells on coverslips were stained for MAP2 after three weeks of differentiation. Neurons were imaged using an Axio Imager Z1 with 568 nm laser light and an Axiocam 506 mono and digitally reconstructed using Neurolucida 360 software (MBF–Bioscience, Williston, ND, USA). When cells were too large for one image, multiple images of these neurons were taken and subsequently stitched together using the stitching plugin of FIJI 2017 software. Only cells that had at least two dendritic branches had been selected for analysis. If the diameter of an extension of the soma was less than 50% of the diameter of the soma, it was an individual primary dendrite. If larger, the extension was evaluated to be made of two different primary dendrites. Axons were not considered in this analysis. For detailed morphological analysis different parameters were chosen to be considered: Branched structure analysis, soma size, number of nodes (dendritic branching points), number of dendrites, dendritic length and the mean length of all cells was conducted. Additionally, convex hull 3D analysis was performed, measuring the area of the dendritic field of a neuron. A net connecting the most distant parts of the neurons dendrites is formed, measuring the volume of the area. Since the pictures obtained were 2D, the convex hull parameter was divided by 2 for all tracings, resulting in a measurement of the surface area and not the full volume. Furthermore, Sholl analysis was performed to investigate dendritic complexity ^82^. Concentric circles are placed in certain, coherently large radii centered at the soma. The distance between two concentric circles forms a shell. The interval chosen for these shells was 10 µm. Within each shell, the number of intersections (the number of dendrites that cross each concentric circle), number of nodes and total dendritic length was calculated.

### Western blot

For Western blot cell lysates were made from iPSC cultures that were 80-90% confluent on a 6-well plate. Medium was always refreshed the day before. Drug treatments were applied as described earlier. To lyse the cells medium was removed and the well was washed with 2 ml ice cold PBS before 100 µl lysis buffer was applied (RIPA buffer supplemented with PhosSTOP (Roche) and protease inhibitors (cOmplete Mini, EDTA free, Roche). Before blotting, the protein concentration was determined by means of a Pierce™ BCA protein assay (Thermo Fisher Scientific). For each sample the same amount of protein between 5 – 7.5 µg was loaded and separated by SDS-PAGE. Depending on the primary antibody separated proteins were transferred to nitrocellulose (BioRad) or, for LC3 probing (1:200; NanoTools 0231-100/LC3-5F10), PVDF membrane (BioRad). Other primary antibodies that were used are: KANSL1 (1:500; Sigma HPA006874), P62 (1:200; Sigma p0067), SOD1 (1:1000; Abcam ab13498), mTOR (1:1000; Cell Signaling 2972), p-mTOR (1:1000; Cell Signaling 2971), GAPDH (1:1000; Cell Signaling 2118), ULK1 (1:1000; Cell Signaling 8054), p-ULK1 (1:1000; Ser757) (1:1000; Cell Signaling 14202), and LAMP1 (1:200; Sigma L1418-200ul). For visualization horseradish peroxidase-conjugated secondary antibodies were used: Goat anti-mouse (1:20000; Jackson ImmunoResearch Laboratories 115-035-062), and Goat anti-rabbit (1:20000; Invitrogen G21234).

### Quantitative polymerase chain reaction

RNA samples were isolated from iPSCs using the NucleoSpin RNA isolation kit (Machery Nagel) according to the manufactures’ instructions. cDNAs were synthesized by iScript cDNA synthesis kit (BioRad) and cleaned up using the Nucleospin Gel and PCR clean-up kit (Machery Nagel). Human specific primers were designed with help of the Primer3plus tool (http://www.bioinformatics.nl/cgi-bin/primer3plus/primer3plus.cgi) (see Supplementary Table 1). PCR reactions were performed in the 7500 Fast Real Time PCR System apparatus (Applied Biosystems) by using GoTaq qPCR master mix 2x with SYBR Green (Promega) according to the manufacturer’s protocol. All samples were analyzed in triplo in the same run and placed in adjacent wells. Reverse transcriptase-negative controls and no template-controls were included in all runs. The arithmetic mean of the C_t_ values of the technical replicates was used for calculations. Relative mRNA expression levels for all genes of interest were calculated using the 2^-ΔΔCt method with standardization to PPIA (Peptidylprolyl Isomerase A) expression level. All expression analyses were done for three different biological replicates in three independent experiments.

### Micro-electrode array recordings and analysis

Recordings of the spontaneous activity of iPSCs-derived neuronal networks were performed at DIV 30 using the 24-well MEA system (Multichannel Systems, MCS GmbH, Reutlingen, Germany). Each well embedded with 12 electrodes, 80 µm in diameter and spaced 300 µm apart). Spontaneous electrophysiological activity of iPSC-derived neuronal network was recorded for 10 minutes. During the recording, the temperature was maintained constant at 37°C, and the evaporation and pH changes of the medium was prevented by inflating a constant, slow flow of humidified gas (5% CO2, 20% O2, 75% N2) onto the MEA. The signal was sampled at 10 kHz, filtered with a high-pass filter (i.e. butterworth, 100 Hz cutoff frequency). The noise threshold was set at ±4.5 standard deviations. Data analysis was performed off-line by using Multiwell Analyzer, software from the Multiwell MEA system that allows the extraction of the spike trains and parameters describing the network activity. The *mean firing rate (MFR)* of the network was calculated by computing the firing rate of each channel averaged among all the active electrodes of the MEA (MFR>0.1 Hz). *Burst detection*. The algorithm defines bursts with a maximum of 50 ms inter spike interval (ISI) to start a burst, and a maximum of 50 ms ISI to end a burst, with a minimum of 100 ms inter burst interval (IBI). The *percentage of random spikes (PRS)* was defined by calculating the percentage of spikes not belonging to a burst for each channel and averaging among all the active electrodes of the MEA. For *Network burst detection* we were looking for sequences of closely-spaced single-channels bursts. A network burst was identified if it involved at least the 80% of the network active channels. *Irregularity* was estimated by computing the CV of the network burst inter burst interval (NIBI), which is the standard deviation divided by the mean of the NIBI. Discriminant function analysis with canonical discriminant functions based on parameters describing neuronal network activity were performed in SPSS.

### Single cell electrophysiology recordings and analysis

For recording spontaneous action potential-evoked postsynaptic currents (sEPSC) we used neurons derived from C_1_, KdVS_1_ and CRISPR_1_ after three weeks of differentiation. Experiments were performed in a recording chamber on the stage of an Olympus BX51WI upright microscope (Olympus Life Science, PA, USA) equipped with infrared differential interference contrast optics, an Olympus LUMPlanFL N 60x water-immersion objective (Olympus Life Science, PA, USA) and kappa MXC 200 camera system (Kappa optronics GmbH, Gleichen, Germany) for visualisation. We performed the recordings of neurons cultured on cover slips under continuous perfusion with oxygenated (95% O2/5% CO2) artificial cerebrospinal fluid (ACSF) (pH 7.4) at 30°C containing 124 mM NaCl, 1.25 mM NaH2PO4, 3 mM KCl, 26 mM NaHCO3, 11 mM Glucose, 2 mM CaCl2, 1 mM MgCl2. Patch pipettes (6-8 MΩ) were pulled from borosilicate glass with filament and fire-polished ends (Science Products GmbH, Hofheim, Germany) using the PMP-102 micropipette puller (MicroData Instrument, NJ, USA). SEPSCs recordings were performed in voltage clamp mode, pipettes were filled with a cesium-based solution containing (in mM) 115 CsMeSO3, 20 CsCl, 10 HEPES, 2.5 MgCl2, 4 Na2-ATP, 0.4 Na3-ATP, 10 Na-phosphocreatine, 0.6 EGTA (adjusted to pH 7.2 and osmolarity 304 mOsmol). Spontaneous action potential-evoked postsynaptic currents (sEPSC) were recorded in ACSF without additional drugs at a holding potential of −60 mV. All recordings were acquired using a Digidata 1140A digitizer and a Multiclamp 700B amplifier (Molecular Devices, Wokingham, United Kingdom), with a sampling rate set at 20 kHz and a lowpass 1kHz filter during recording. Recordings were not analyzed if series resistance was above 20 MΩ or when the recording reached below a 10:0 ratio of membrane resistance to series resistance. SEPSCs were analyzed using MiniAnalysis (Synaptosoft Inc). For the traces that contained strongly accumulated sEPSCs mediated by presynaptic synchronized bursts of action potentials, these bursts of sEPSCs were counted by visual inspection.

### Statistics

The statistical analysis of the data was performed using GraphPad Prism 8 (GraphPad Software, Inc., CA, USA). We first determined whether data was normally distributed. We tested statistical significance for different experimental conditions by one-way ANOVA or two-way ANOVA when different cell-lines and drug-treated samples were included. Individual samples were compared using Sidak’s or Dunnett’s multiple comparisons test. When only two conditions were compared, we used unpaired t test. When data was not normally distributed we applied Kruskal-Wallis test combined with Dunn’s or Sidak’s multiple comparison test. Results with *P* values lower than 0.05 were considered as significantly different (*), *P* <0.01 (**), *P* <0.001 (***), *P* <0.0001 (****). Data is shown in violin plots or as mean and standard error of the mean (SE) in bar diagrams. Details about statistics are reported in the Supplementary Table S3.

## Author contributions

K.L. N.N.K., D.K., B.d.V conceived and supervised the study. K.L. and N.N.K. designed all the experiments. K.L., E.L., A.V., T.K.G., L.D, E.U, A.O, C.S. and N.N.K. performed all the experiments. G.T., M.G. provided resources. D.S., M.F., K.L. and N.N.K. performed data analysis. K.L and N.N.K. wrote the paper. D.S. D.K., B.d.V. edited the paper.

