## supplemental data for "KANSL1 Deficiency Causes Neuronal Dysfunction by Oxidative Stress-Induced Autophagy"

Radboudumc

Department of Human Genetics

Geert Grooteplein 10

P.O.Box 9101

Nijmegen 6500HB

the Netherlands

**List Supplementary Material**

- Supplementary Figure 1 (Characterization of control and KdVS iPSCs)
- Supplementary Figure 2 (Autophagy phenotype in control and KdVS iPSCs)
- Supplementary Figure 3 (Mitochondrial function in control and KdVS iPSCs)
- Supplementary Figure 4 (iNeuron morphology of control and KdVS cells)
- Supplementary Figure 5 (Cell Density and autophagy phenotype in control and KdVS iNeurons)
- Supplementary Figure 6 (Reclassification of group membership based on MEA parameters)
- Supplementary Figure 7 (Oxidative stress induced autophagy in KdVS)
- Supplementary Table 1 (List of qPCR primers)
- Supplementary Table 2 (List of vectors)

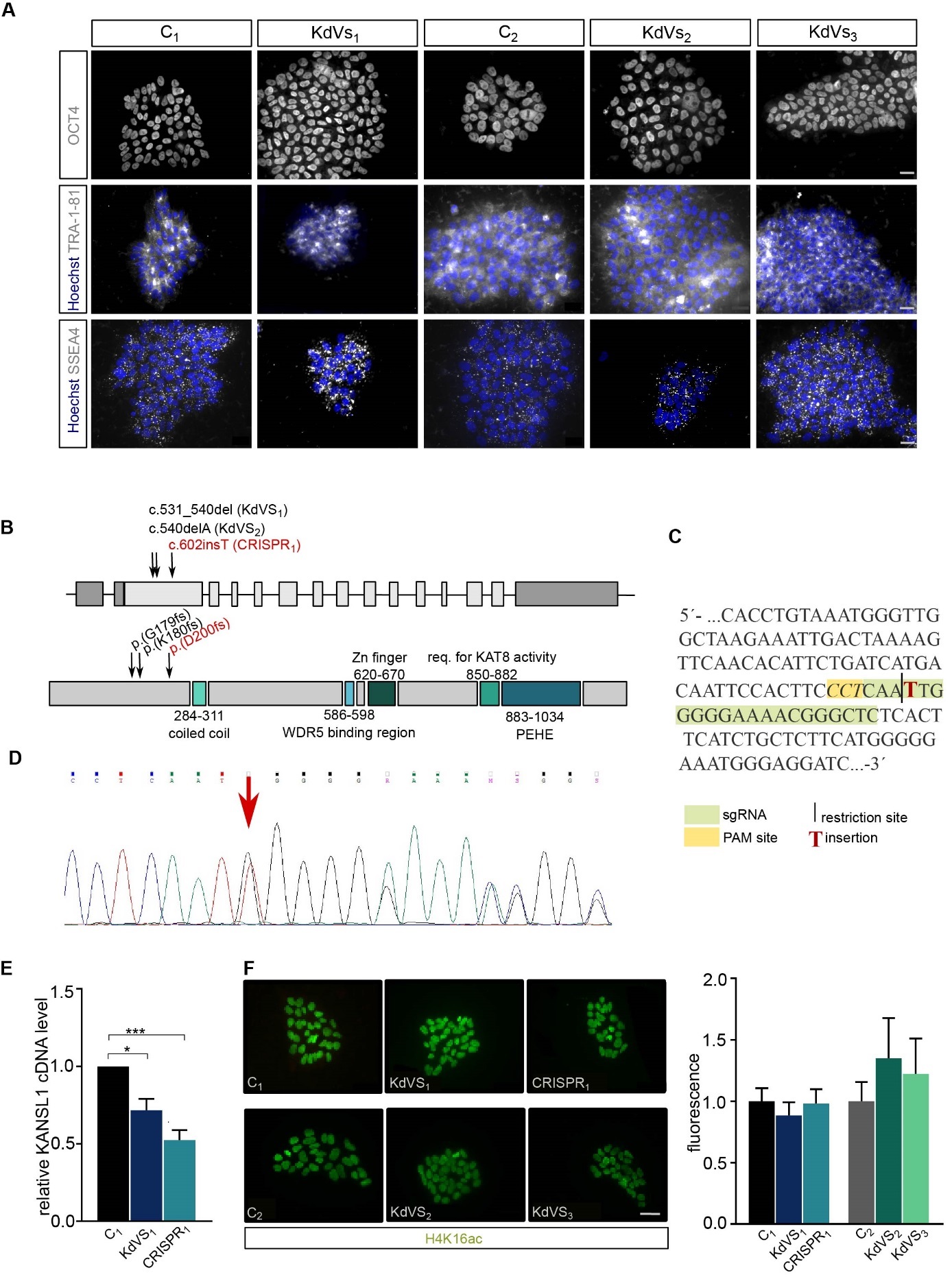

***Supplementary Figure 1. iPSC characterization.* A)** Representative images for pluripotency marker detection in iPSC lines. Scale bar = 20 µm. **B)** Schematic presentation of the KANSL1 gene and protein. Mutations in KANSL1 deficient lines are marked. **C)** Nucleotide sequence of KANSL1 gene. Exon 2 that was used for sgRNA design and CRISPR/ Cas9 editing. **D)** Sanger sequencing result for CRISPR_1_ indicating a one- nucleotide insertion in one allele at the targeted restriction site. **E)** q-PCR result for KANSL1 to prove lower expression in KdVS_1_ patient derived iPSC line and CRISPR_1_ when compared to the respective control line C_1_. n=6 for all samples. **F)** Representative images for iPSC colonies stained for H4K16ac and H4K16ac quantification. n = 13 for C_1_, KdVS_1_ and CRISPR_1_; n = 7 for C_2_, KdVS_2_ and KdVS_3_. Fluorescence for KANSL1 deficient lines was normalized to the respective control lines. Scale bar = 50µm.Ordinary one-way ANOVA and Sidak’s multiple comparison test were used to test for statistically significant difference. **P* < 0.05, ***P* < 0.01, ****P* < 0.005, *****P* < 0.0001.

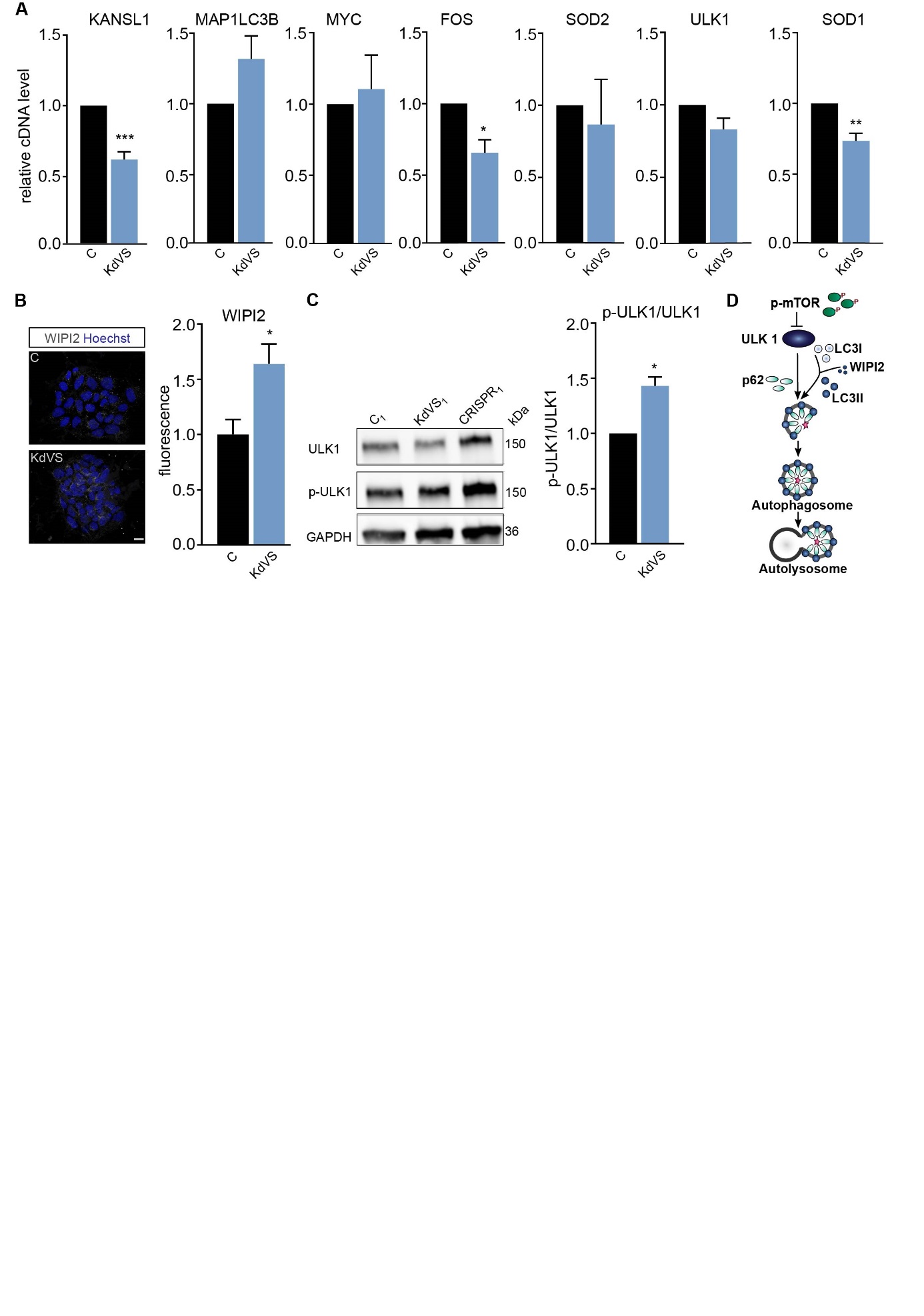
**Supplementary Figure 2. Autophagy Phenotype in iPSCs.** **A)** qPCR results for several H4K16ac regulated ATGs and KANSL1. cDNA levels in KdVS patient derived lines were always compared to levels in control iPSCs. n = 3. **B)** Representative images and signal quantification of iPSC colonies stained for autophagosome marker WIPI2. KdVS patient derived cells were compared to control iPSCs. n= 5 for control; n=12 for KdVS. Fluorescence was normalized to mean fluorescence of the respective control per experiment. Scale bar= 20 µm. **C)** Representative Western Blots and quantification for ULK1 phosphorylation (Ser757). P-ULK1/ ULK1 ratio was normalized to control per experiment. n = 4 for control; n = 8 for KdVS. **D)** Schematic representation of mTOR dependent autophagy activation and autophagosome formation. Decreased mTOR phosphorylation will enable ULK complex activation which then induces phagophore formation. LC3I is lipidated to LC3II with help of WIPI2. P62 serves as adaptor protein between LC3II and autophagosomal cargos. Upon completion autophagosomes fuse with lysosomes to form autolysosomes and subsequently degrade cargos including p62. Statistically significant differences were determined by means of unpaired t test. *P < 0.05, **P < 0.01, ***P < 0.005, ****P < 0.0001.

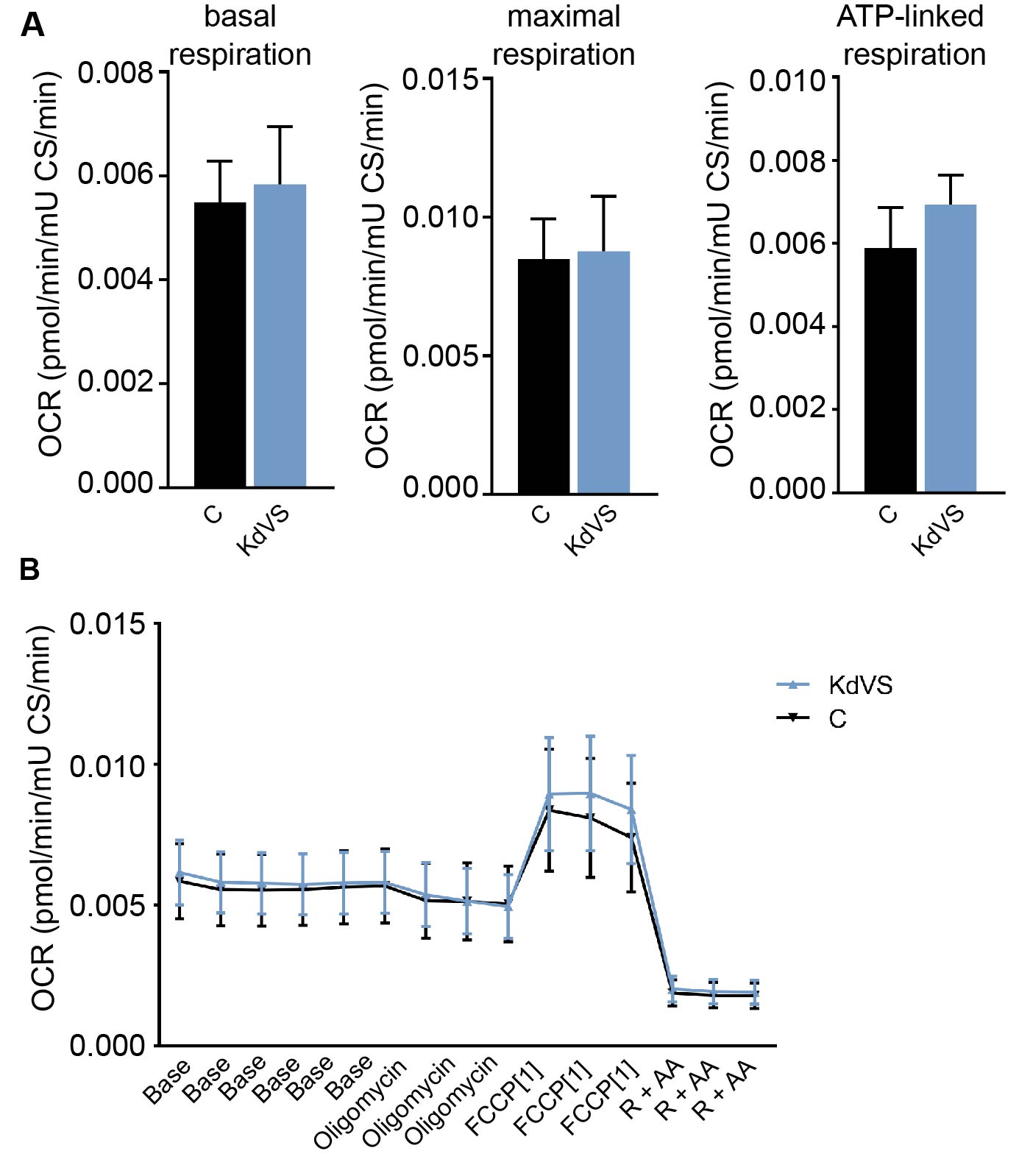

***Supplementary Figure 3. Mitochondrial function.*** **A)** Seahorse assay results for KdVS compared to control iPSCs showing basal, maximal, and ATP-linked respiration. **B)** We used oxygen consumption rate (OCR) as a measure of mitochondrial respiration and normalized to oxaloacetate-induced citrate synthase activity. n = 15 per condition for each line. Statistically significant differences were determined by means of t test. **P* < 0.05, ***P* < 0.01, ****P* < 0.005, *****P* < 0.0001.

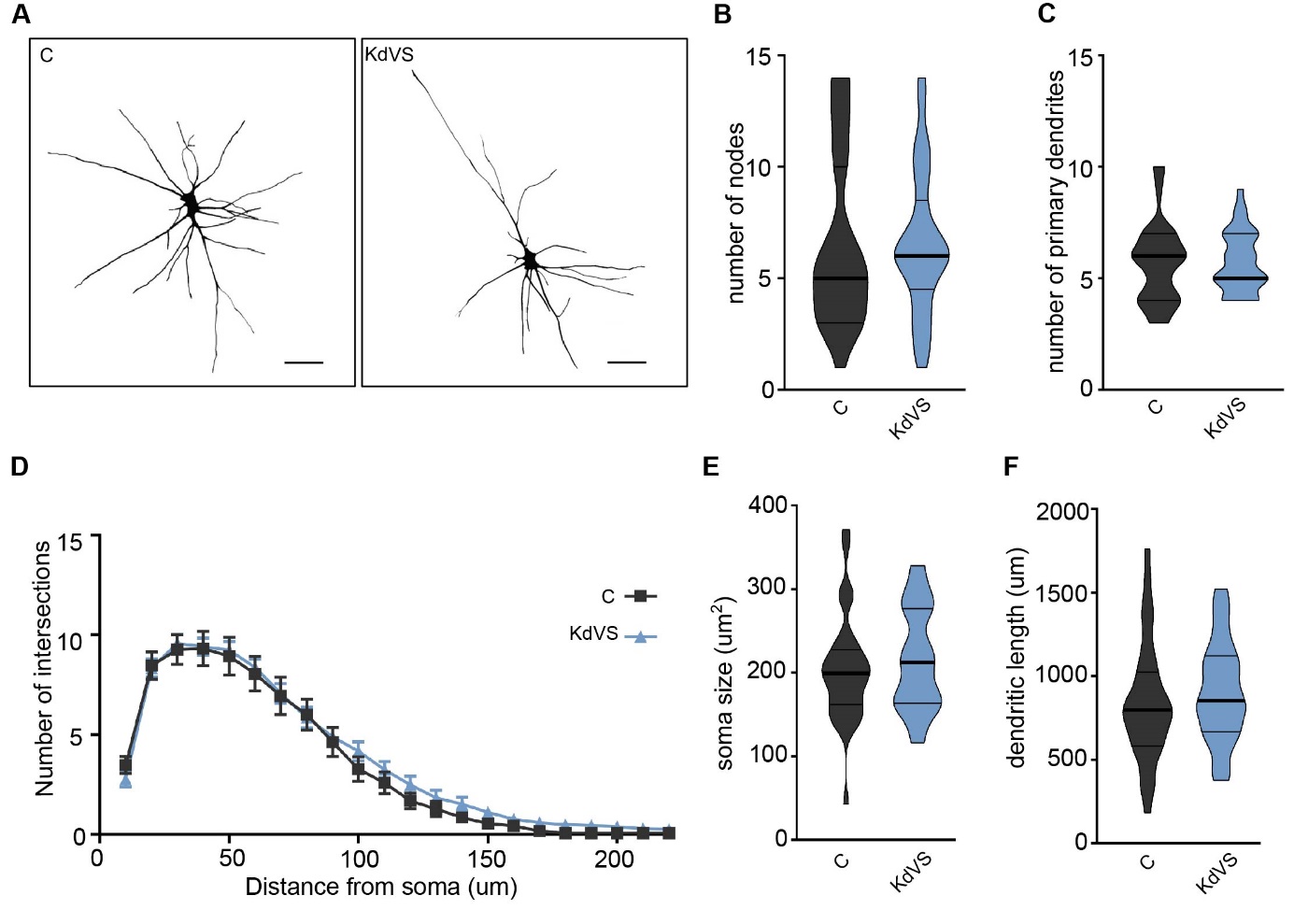
***Supplementary Figure 4. Neuronal Morphology.* A)** Representative images of reconstructed neurons derived of KdVS_1_ and C_1_ iPSCs, respectively. Scale bar = 50µm. **B)-F)** Detailed morphological analysis revealed no significant differences in morphology between the different lines. n = 33 for control and n = 29 for KdVS. Statistical differences were determined by t test. **P* < 0.05, ***P* < 0.01, ****P* < 0.005, *****P* < 0.0001.

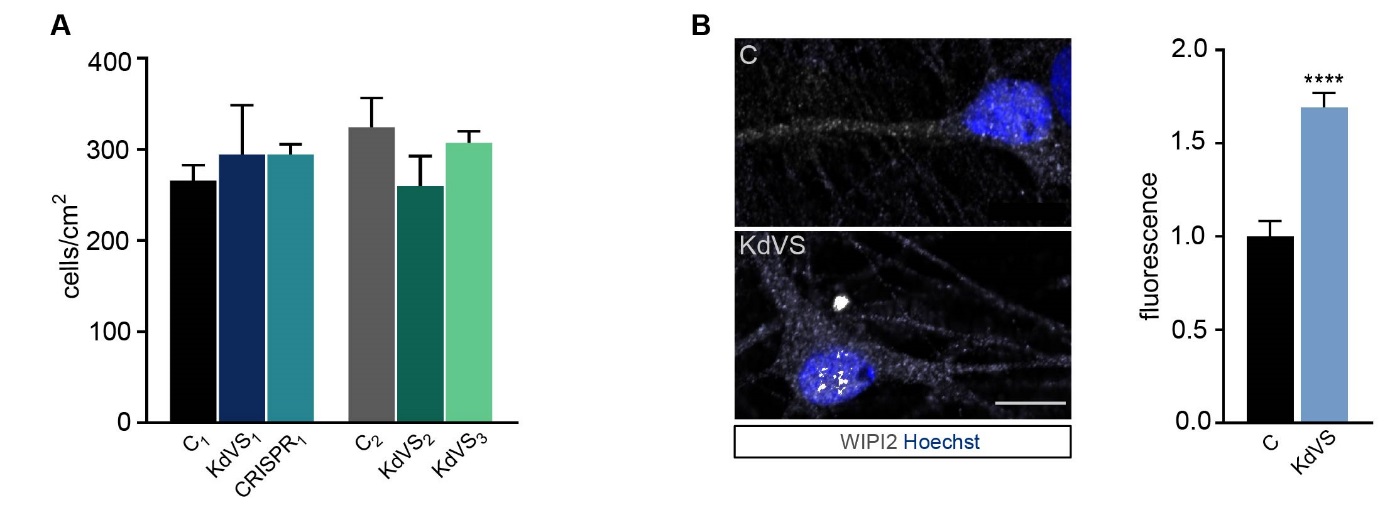
 ***Supplementary Figure 5. Neuronal Cell Density and Autophagy phenotype.* A)** Cell counts of fixed and stained iNeurons on c/s at DIV21 to ensure comparable cell densities between the lines and experiments. Ordinary one-way ANOVA was used for statistical analysis. **B)** Representative images and quantification for WIPI2 at DIV21. n = 25 for control; n = 31 for KdVS. Fluorescence was normalized to control. Scale bar= 20 µm. Unpaired t test was used to test for statistically significant differences. **P* < 0.05, ***P* < 0.01, ****P* < 0.005, *****P* < 0.0001.

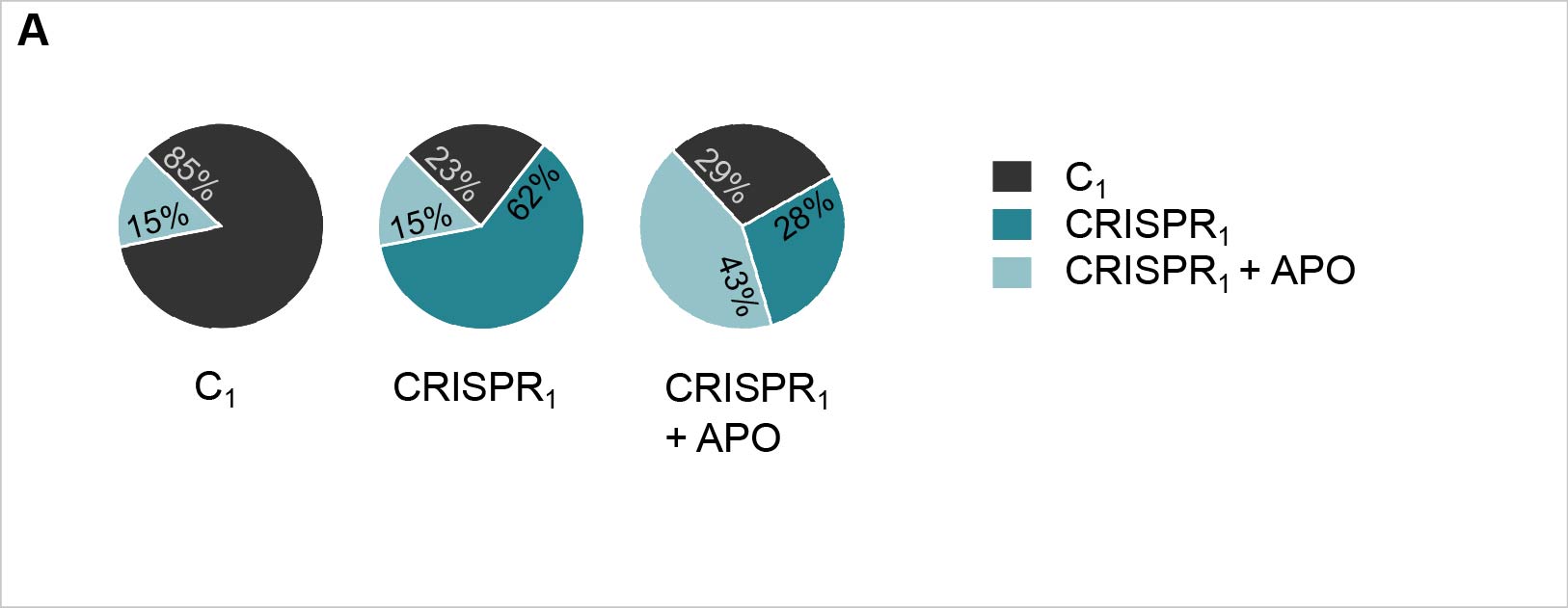
***Supplementary Figure 6. Reclassification of group membership.* A)** Pie charts visualize accuracy of discriminant analyses functions by showing the relative distribution of lines per a-priory group after reverse testing for group identity.

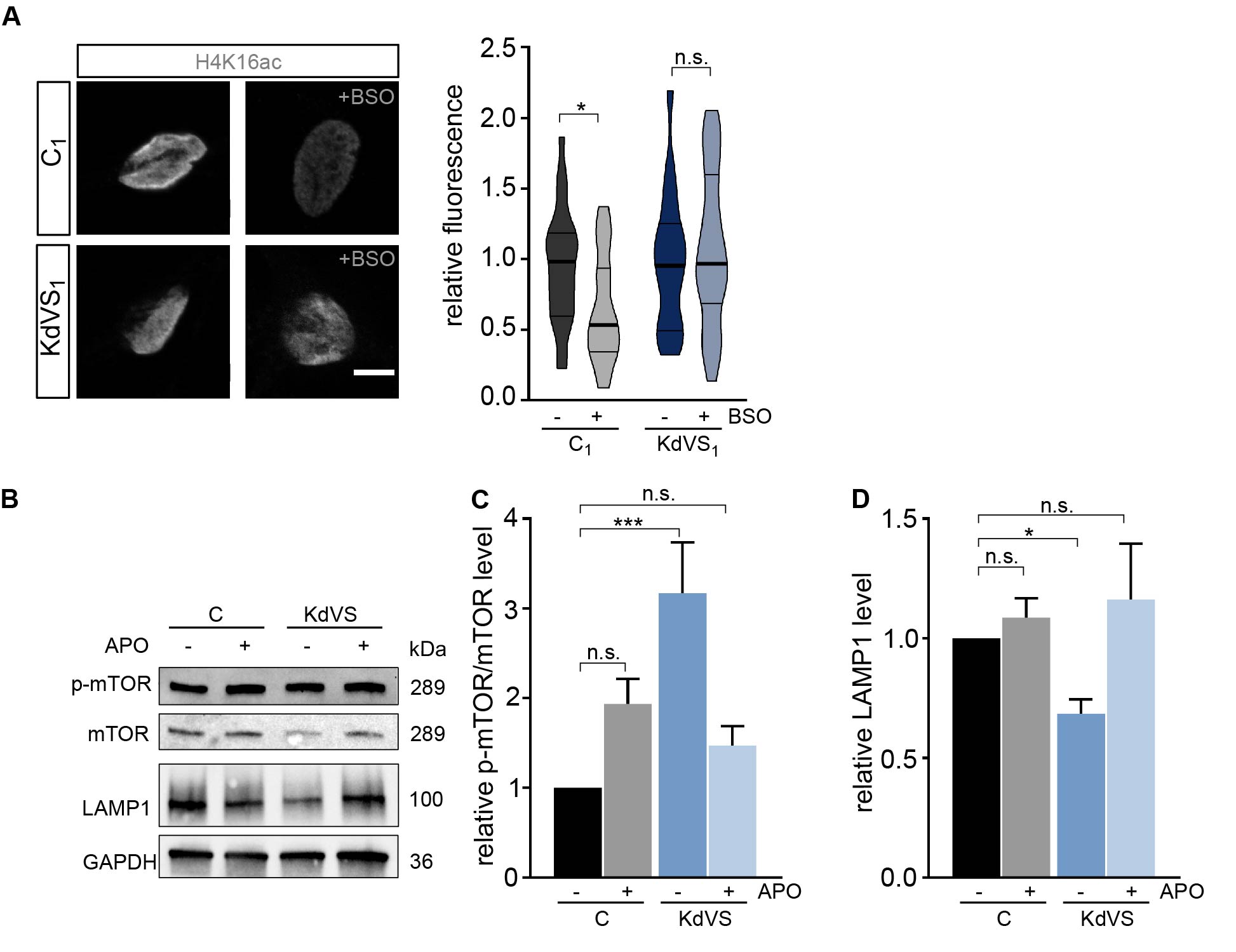
***Supplementary Figure 7. Oxidative stress causes KdVS phenotype.* A)** Representative images of iNeuron nuclei stained for H4K16ac with and w/o BSO driven autophagy induction for C_1_ and KdVS_1_ and fluorescence quantification. All fluorescence values were normalized to the untreated control. Scale bar = 20 µm. n = 21 for C_1_; n = 26 for C_1_+BSO; n = 17 for KdVS_1_; n = 18 for KdVS_1_+BSO. Ordinary one-way ANOVA was used for statistical analysis. **B-D)** Representative Western blots for p-mTOR, mTOR, and LAMP1. iPSCs were either untreated or treated o/n with 100 µM APO. p-mTOR/mTOR an LAMP1 levels were quantified relative to untreated control. n = 5 for C_1_; n = 5 for C_1_+APO; n = 7 for KdVS; n = 7 for KdVS+APO. Statistically significant differences were determined by means of Kruskal Wallis and Dunn’s multiple comparison test. **P* < 0.05, ***P* < 0.01, ****P* < 0.005, *****P* < 0.0001.

***Supplementary Table 1. List of qPCR primers.***

| Gene | Direction | Sequence |
| --- | --- | --- |
| KANSL1 | forward | ATCCTCCACACAGTCCCTTG |
|  | reverse | CCCCTTCTCCTCCTTACTGG |
| MAP1LC3B | forward | GAGAAGCAGCTTCCTGTTCTGG |
|  | reverse | GTGTCCGTTCACCAACAGGAAG |
| ULK1 | forward | GTTCCAAACACCTCGGTCCT |
|  | reverse | GCTCAGGGATGGTTCCAACT |
| SOD1 | forward | TGAAGAGAGGCATGTTGGAGAC |
|  | reverse | CAAGCCAAACGACTTCCAGC |
| SOD2 | forward | CTGCTCCCCGCGCTTT |
|  | reverse | GCTGGTGCCGCACACT |
| FOS | forward | GCGTTGTGAAGACCATGACAG |
|  | reverse | TCTAGTTGGTCTGTCTCCGCT |
| MYC | forward | CAGCGACTCTGAGGAGGAAC |
|  | reverse | GCTGCGTAGTTGTGCTGATG |

***Supplementary Table 2. List of vectors.***

| Plasmid | Source | Experiment |
| --- | --- | --- |
| pLVX-EF1α-(Tet-On-Advanced)-IRES-G418(R) | J. Ladewig | Generating Ngn2/rTTA positive iPSC lines |
| pLVX-(TRE-thight)-(MOUSE)Ngn2-PGK-Puromycin(R) | J. Ladewig | Generating Ngn2/rTTA positive iPSC lines |
| psPAX2 lentiviral packaging vector | Addgene #12260 | Lentiviral preparations |
| pMD2.G lentiviral packaging vector | Addgene #12259 | Lentiviral preparations |
| FUW mCherry-GFP-LC3 | Addgene  #110060 | Autophagosome/ synapsin co-localization |
